# Analysis of *Phycodnaviridae* Tetrapyrrole Enzymes: Subcellular Localization and Functional Divergence from Putative Host Homologs

**DOI:** 10.64898/2026.08.07.743453

**Authors:** Steven Zehnacker, Stefano Caffarri, Guillaume Blanc, Xenie Johnson, Marina I. Siponen

**Affiliations:** Photosynthesis and Environment team and ProteinTec, Institute of Bioscience and Biotechnology of Aix-Marseille University, CEA/CNRS/AMU, France; Luminy Genetics and Biophysics of Plants Team, Institute of Bioscience and Biotechnology of Aix-Marseille University, CEA/CNRS/AMU, France; Environmental Microbiology and Biotechnology, Mediterranean Institute of Oceanology, CNRS, Marseille, France

**Keywords:** Phycodnaviridae, Auxiliary metabolic genes, Protein targeting, heme catabolism, Bilins, Heme oxygenase (HMOX1), Phycocyanobilin: Ferredoxin oxidoreductase (PcyA)

## Abstract

**Rationale:** Recent viral metagenomic studies have revealed a broad repertoire of enzyme encoding genes in *Phycodnaviridae*, whose contributions to viral replication and host infection remain poorly understood. These genes may expand the metabolic repertoire of viruses during infection of green algal hosts. Because neither the roles of these proteins during infection nor their subcellular localization have been experimentally characterized, comparative sequence, structural, and biochemical analyses done *in silico* can generate testable hypotheses on structure, function and localization.

**Methods:** In a recent viral metagenomic dataset, we identified viral homologues of two eukaryotic enzymes involved in bilin biosynthesis: heme oxygenase-1 (vHMOX1) and Phycocyanobilin:Ferredoxin oxidoreductase (vPcyA). Viral and algal homologues were compared through sequence analyses and AlphaFold3 structural predictions. Predicted biochemical properties were analyzed for their compatibility with subcellular compartments. Active site architecture and putative substrate binding were compared between viral and algal proteins using homologous structures deposited in the Protein Data Bank (PDB).

**Results:** Viral HMOX1 and PcyA sequences are truncated compared to algal homologues, lacking the N-terminal extension associated with chloroplastic targeting. However biochemical properties, including isoelectric point and surface charge distribution, are compatible with localization in chloroplast. Structural comparisons reveal modifications in the viral HMOX1 active sites, including altered substrate binding and substitutions of key residues, likely indicating modified heme-binding properties. In addition to these modifications, chloroplastic ferredoxin interaction predictions show compatibility for use of this electron donor. In contrast, vPcyA models show no significant differences with their algal counterparts.

**Key findings:** Our bioinformatic analyses expand the repertoire of auxiliary metabolic genes in Phycodnaviridae by identifying a conserved heme degradation pathway, non-canonical vHMOX1/PcyA targeting and structural rearrangements surrounding the catalytic sites of viral HMOX1.

## Introduction

Viruses infecting photosynthetic microorganisms are major drivers of marine ecosystem dynamics, influencing primary production, nutrient cycling, and microbial community structure (Fuhrman 1999; Suttle 2005, 2007; Wileman *et al*. 2016). Over the past decade, advances in viral metagenomics have revealed that many marine viruses harbor auxiliary metabolic genes (AMGs), encoding proteins involved in host metabolic pathways (Brahim Belhaouari *et al*. 2022). These genes are thought to modulate host physiology during infection, thereby optimizing viral replication. This concept has been documented in bacteriophages infecting cyanobacteria, where AMGs involved in photosynthesis, carbon metabolism (Lindell *et al*. 2005; Thompson *et al*. 2011; Ledermann *et al*. 2016; Wegner *et al*. 2024), and stress responses have been functionally characterized. In contrast to bacteriophages, AMGs in viruses infecting eukaryotic phototrophs, such as members of the *Phycodnaviridae* and related lineages, remain comparatively unexplored. To date, the only targets studied have been linked to metabolism and ROS management and are postulated to optimize the efficiency of viral replication during different stages of the infection process (Rosenwasser *et al*. 2016). Understanding the functional utility of AMGs is particularly complex and challenging in eukaryotic hosts, where metabolic pathways are highly compartmentalized, notably within plastids.

A recent study by Chase *et al.,* 2024, (Chase *et al*. 2024) reported the discovery of large viruses putatively associated with the green algae *Picochlorum* that encode homologues of enzymes involved in the chloroplastic bilin biosynthesis pathway: Heme oxygenase 1 (HMOX1) and Phycocyanobilin:Ferredoxin Oxidoreductase (PcyA). This finding is unprecedented, as viral-encoded components of tetrapyrrole metabolism had not previously been described in eukaryotic oceanic viruses. In eukaryotic photosynthetic organisms, this pathway is tightly regulated and the product, phycocyanobilin, plays multiple essential roles in photosynthesis depending on the species including: light capture (phycobilisomes), light sensing (phytochromes) and chlorophyll biosynthesis (stabilization and protection of Mg-Chelatase) (Duanmu *et al*. 2013; Wittkopp *et al*. 2017; Zhang *et al*. 2021; Wang *et al*. 2026). The presence of genes for these enzymes in viral genomes infecting green algae raises fundamental questions regarding their function and intracellular localization.

A key aspect of virus infection in eukaryotic cells is the formation of viral replication compartments, or viral factories (VF), which arise from extensive remodeling of the host cytoplasm (Netherton *et al*. 2007; Netherton and Wileman 2011; Fernández de Castro *et al*. 2021). These structures concentrate viral and host components and create a distinct biochemical environment that supports viral replication. Viral-encoded metabolic enzymes could be in host organelles, such as the chloroplast, or instead function within the cytoplasm, in the VF. Notably, homologous enzymes of HMOX1 and PcyA in green algae typically possess N-terminal chloroplast transit peptides and are imported into plastids, where tetrapyrrole metabolism is located (Duanmu *et al*. 2013). Furthermore, synthesis of both biliverdin and phycocyanobilin requires electrons from ferredoxin which is found only in chloroplasts or mitochondria. Whether viral homologs retain such targeting or have evolved to operate in alternative compartments remains an open question.

In the absence of experimentally tractable host–virus systems, *in silico* approaches provide a powerful framework to address these questions. Recent advances in protein structure prediction using artificial intelligence, such as AlphaFold3 (Abramson *et al*. 2024), enable detailed analysis of predicted protein architecture, surface properties (*e.g*., surface charge distribution, hydrophobicity), and potential substrate and/or protein interactions. Combined with sequence-based analyses of N-terminal targeting signals and biochemical properties (*e.g*., isoelectric point), these approaches can potentially allow inference of protein localization and their functional adaptation. Here, we investigate viral homologs of heme oxygenase 1 (vHMOX1) and Phycocyanobilin: Ferredoxin oxidoreductase (vPcyA) identified in algal-associated viral genomes. By combining sequence analysis, prediction of N-terminal targeting features, and structural modeling, we discuss where these viral enzymes are likely to be targeted. We further characterize their biochemical and structural properties to infer potential functional adaptations, providing insights into how viruses may reprogram host tetrapyrrole metabolism during infection.

## Materials and Methods

### Identification of HMOX1 and PcyA homologues

Viral homologues of heme oxygenase (HMOX1) and Phycocyanobilin: Ferredoxin oxidoreductase (PcyA) were identified among proteins predicted from viral metagenomic assemblies generated by Chase *et al*. 2024, using PSI-BLAST. HMOX1 and PcyA proteins from *Picochlorum* sp. BPE23 (GenBank accessions KAI8104144.1 and KAI8100569.1, respectively) were used as initial queries against the predicted metagenomic protein dataset. Searches were run for three iterations with an inclusion E-value threshold of 1 × 10⁻⁵, allowing the construction of position-specific scoring matrices to detect divergent homologues. Hits with E-values below 1 × 10⁻⁵ were retained. Candidate sequences were subsequently queried against the Swiss-Prot database to confirm their assignment as HMOX1 or PcyA homologues before downstream analyses.

### N-terminal sequences analyses and gene annotation

Multiple sequence alignments of HMOX1 and PcyA homologs (Fig. 1A and 1B) were performed. Homologous sequences were identified using BLAST searches (AltschuP *et al*. 1990). Multiple sequence alignments were generated using MUSCLE or MAFFT with default parameters (Edgar 2004) and manually curated to remove poorly aligned regions, particularly within variable N-terminal extensions. and manually curated to remove poorly aligned regions, particularly within variable N-terminal extensions. Complete protein sequence alignments (Fig. S1 and Fig. S2), including protein identification numbers, are provided in the Supplementary Data (Table S1A and Table S1B). Additional *Picochlorum* sequences were retrieved from available genome assemblies to increase the number of potential host homologues. Specifically, sequences from *Picochlorum celeri* were derived from the GenBank genome assembly GCA_010909725.1; *Picochlorum. oklahomense* sequences were derived from the genome assembly available at 10.1093/molbev/msy167; SENEW3 sequences were derived from GCA_034226945.1; *Picochlorum costavermella* sequences were derived from GCA_011316045.1; and *Picochlorum atomus* sequences were obtained from the OneKP project database (ID: scaffold-JQFK-2002893- *Picochlorum*_*atomus*). Sequence alignments were visualized and annotated using Geneious. The presence of a transit peptide was predicted using TargetP2.0 (Almagro Armenteros *et al*. 2019).

**Figure 1:**
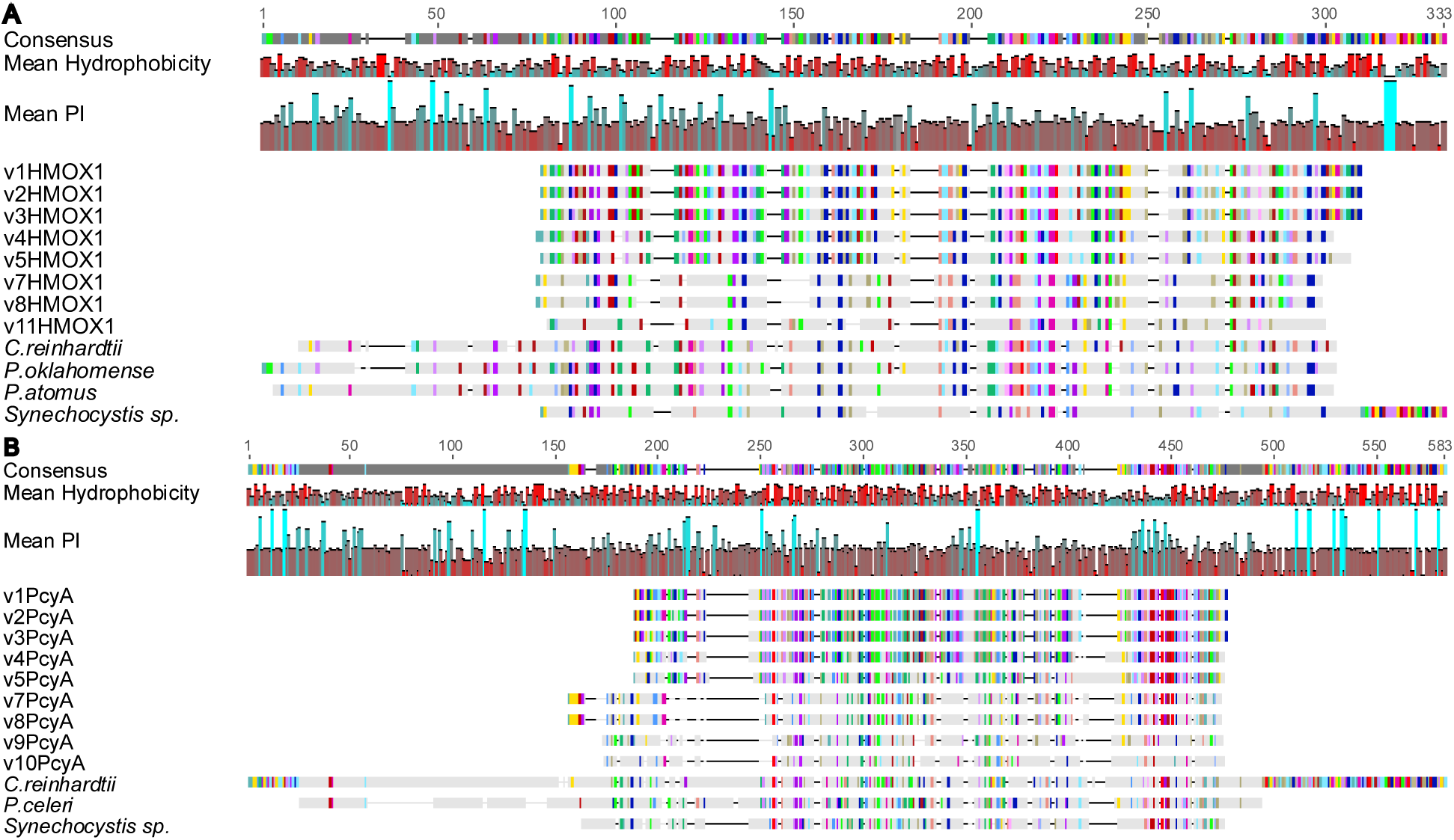
Schematic representation of the multiple-sequence alignment of (A) 9 vHMOX1 sequences and (B) 9 vPCYA sequences identified in the viral metagenomic datasets of Chase *et al*. (2024). Viral sequences were aligned with representative *Picochlorum* homologues and functionally characterized proteins of the model algae *Chlamydomonas reinhardtii* and the model cyanobacteria *Synechocystis PCC6803*. Blocks of color represent conserved residues. The consensus sequence is shown for numbering amino acids and for the identification of the catalytic residues of HMOX1 shown underneath as dark blue bars for 1 residue or arrows that represent 2 conserved residues; mean hydrophobicity reports red as the most hydrophobic residues through to purple and blue as the most hydrophilic residues; Mean pI, stands for Isoelectric point and refers to the mean pH for the amino acid at that position for which a molecule carries no net electrical charge. The lowest pI value is ranked 0 (red) and the highest pI value is ranked 1 (blue) all other residues are ranked accordingly and an average pI along the sequence can be calculated.

### Gene annotation and Alphafold3 modelling

Protein domains of the vHMOX1 and vPcyA proteins were identified using the NCBI CD-SEARCH tool (Marchler-Bauer and Bryant 2004; Lu *et al*. 2020). The 3D modeling of algal and viral HMOX1 and PcyA were done using AlphaFold 3 server (Abramson *et al*. 2024). Key residues were visualized within the viral sequences using Jalview software (Waterhouse *et al*. 2009). After structure prediction, whole models were visualized by ChimeraX 1.8 (Pettersen *et al*. 2021), and subsequently structurally aligned with their closest crystallographic homologues, as identified by the DALI server (http://ekhidna2.biocenter.helsinki.fi/dali/) (Holm *et al*. 2023) and by the SWISS-MODEL database (Waterhouse *et al*. 2018) : HMOX1 from *Glycine max* (PDB: 7CKA, (Tohda *et al*. 2021) and PcyA from *Synechocystis* (PDB: 2DE1, (Hagiwara *et al*. 2010)). The different predicted or experimental models were analyzed using Proteins*Plus* (Ehrt *et al*. 2025) for investigation into the heme ligand binding pocket in the reference *Glycine max* structure (PDB:7CKA), and predicted PicoHMOX1, and v1HMOX1 as the representation of group I enzymes and v7HMOX1 as the representative of group II enzymes. The KAI8100517.1 protein sequence from *Picochlorum sp.* BPE23 was used for ferredoxin modeling, it was identified as the closest *Picochlorum sp.* homolog to the published structure *Mays* (PDB: 9A8X).

## Results

### vHMOX1 and vPCYA sequence analysis

The viral metagenome putatively associated with *Picochlorum* was previously reported to encode the auxiliary metabolic genes *vHMOX1* and *vPCYA* (Chase *et al*., 2024). Here, we have characterized these viral proteins using comparative sequence and structural analyses. Re-examination of the metagenomics dataset identified 12 *vHMOX1* and 10 *vPCYA* genes. The main features of the encoded proteins are summarized in Table 1.

**Table 1:**
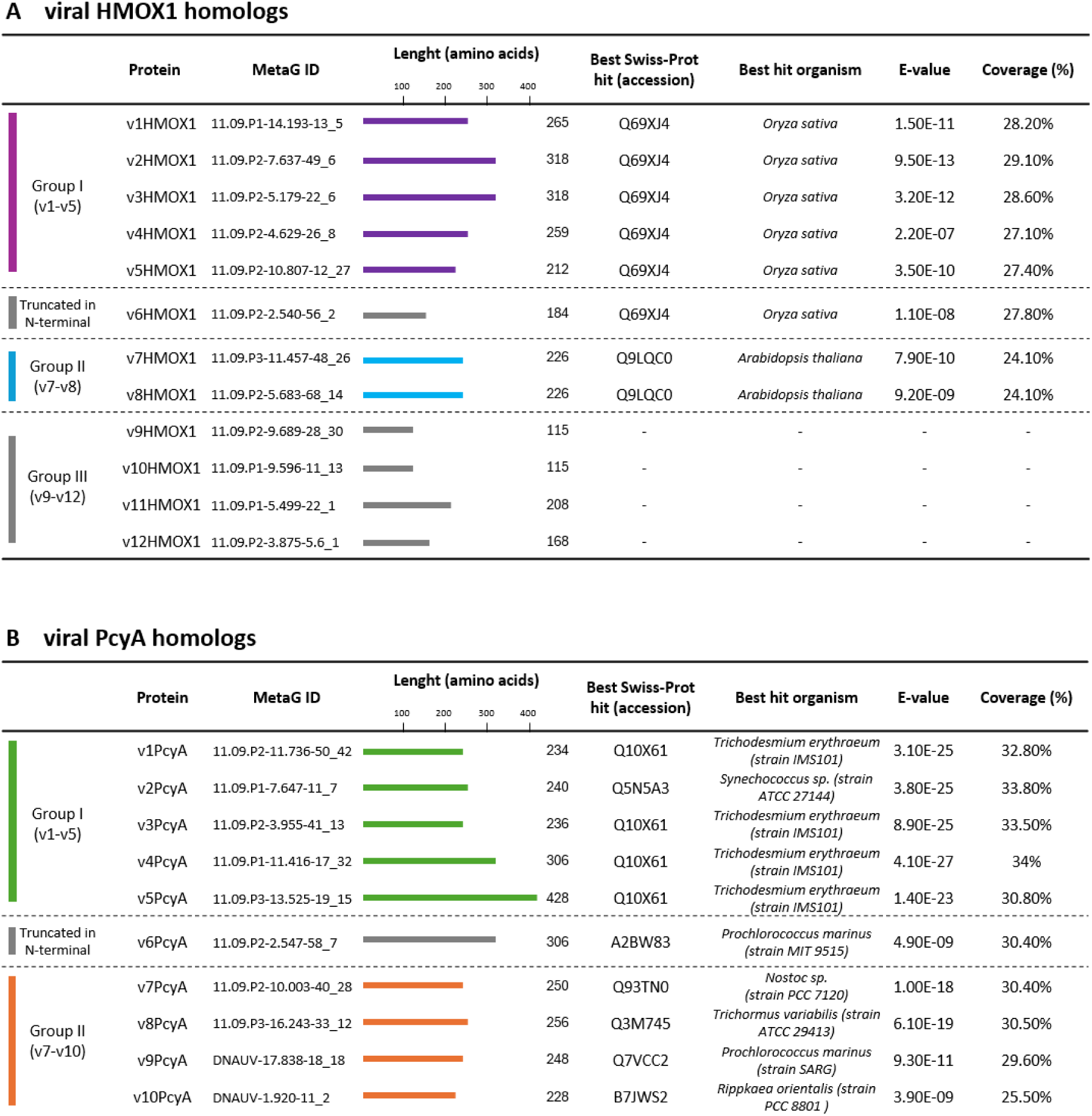
Identification and annotation of viral HMOX1 and PcyA homologs. A: Characteristics of the 12 viral HMOX1 homologs identified in the metagenomic dataset. Proteins are grouped according to their sequence relationships, with Groups I (v1–v5), II (v7–v8) and III (v9-v12 that are highly divergent and did not yield significant Swiss-Prot hits). **B**: Characteristics of the 10 viral PcyA homologs identified in the dataset, separated in Group I (v1-v5) and group II (v7-v10) according to sequence relationships. For each protein, the amino acid length, best Swiss-Prot hit, corresponding organism, BLASTP E-value, and query coverage are indicated. Horizontal bars represent protein length and are scaled according to the amino acid length. Swiss-Prot accessions and best-hit organisms correspond to the top BLASTP matches. Coverage represents the proportion of the viral protein sequence covered by the best hit.

Both the vHMOX1 and vPcyA sequences are diverse in both composition and length (115 amino acids to 318 amino acids for vHMOX1 and 234 amino acids - 428 amino acids for vPcyA sequences). v9-v12 HMOX1 that were initially identified as HMOX1 homologs by PSI blast using *Picochlorum* BP23 homologs did not find any hit in Swiss-Prot database whereas v1-v5 and v7- v8 HMOX1 have respectively HMOX1 from *Oryza sativa* and *Arabidopsis thaliana* as best hits. Because v6 is truncated in N-terminal and v9, v10 and v12 sequences are also shorter and highly divergent compared to other viral HMOX1, we decided to focus our analysis on the group I (v1- v5), group II (v7-v8) and group III (v11) HMOX1 sequences (Table 1A). Sequence annotation shows that v11HMOX1 from group III lacks many of the conserved residues identified as functional for BV production and may also be truncated (Fig. S3). The vPcyA proteins formed two groups: group I (v1-v5) and group II (v7-v10) (Table 1B). All vPcyA find their best hit in Swiss-Prot data base in different cyanobacteria strains. Because v6PcyA is identical to v5PcyA but has a truncated C-term part, we exclude this sequence. Sequence annotations confirm that neither group contains proteins with missing known functional residues (Fig.S4).

Compared to homologous sequences from green algae, the vHMOX1 sequences appear truncated (Fig. 1A, and for the complete alignments see Fig. S1), initiating close to the conserved catalytic domain and lacking the typical N-terminal extension. In this respect, they resemble the HMOX1 protein of the cyanobacterium *Synechocystis*. The *Chlamydomonas reinhardtii* HMOX1 and PcyA proteins were shown to be chloroplast localized in Duanmu *et al*. 2013 (Duanmu *et al*. 2013) and the proteins from different *Picochlorum* shown in Fig. 1A and Fig. B have long N-terminal extensions with predicted chloroplast transit peptides using TargetP2.0 (Fig. 1A, Fig. 1B, Table S2A and Table S2B). Consistently, no chloroplast transit peptide is predicted from the first methionine of the viral sequences, and the N-terminal regions do not show a strong enrichment in Ser/Thr residues, which are commonly observed in plastid transit peptides (Tardif *et al*. 2012). Altogether, these features argue against chloroplastic targeting via a cleavable transit peptide into the chloroplast. The mean isoelectric point (pI) averaged over the region occupied by vHMOX1 is 6.05 and for vPcyA is 5.92 showing no particular preference for the more alkaline stromal environment. The viral proteins, however, do not strongly differ from the algal proteins having the same acidic pI. The mean hydrophobicity is low and randomized in both cases suggesting that these are highly soluble proteins.

### Overall vHMOX1 structural analysis

Because compartmentalization may contribute to the functional diversification of structurally similar HMOX1 enzymes, structural analysis can provide valuable clues to their potential localization and function. Given the apparent absence of canonical plastid-targeting signals in viral HMOX1, we therefore examined their predicted structures to guide hypotheses for future localization studies.

To study the structural conservation of vHMOX1 enzymes, the AlphaFold3 server was used to generate predicted models of these enzymes with or without heme as a cofactor. As expected, the viral enzymes reveal the same canonical fold as other HMOX1 enzymes with the closest homologue identified using the Dali server (http://ekhidna2.biocenter.helsinki.fi/dali/) (Holm *et al*. 2023) as the Glycine max HMOX1 (GmHMOX1) structure (PDB ID:7CKA). Viral HMOX1 retained the canonical HMOX1 fold, with high prediction confidence (pLDDT > 90) and close structural similarity to the *Picochlorum* HMOX1 structures (Fig. 2, Fig. S3, Fig. S5 and Table S3A).

**Figure 2:**
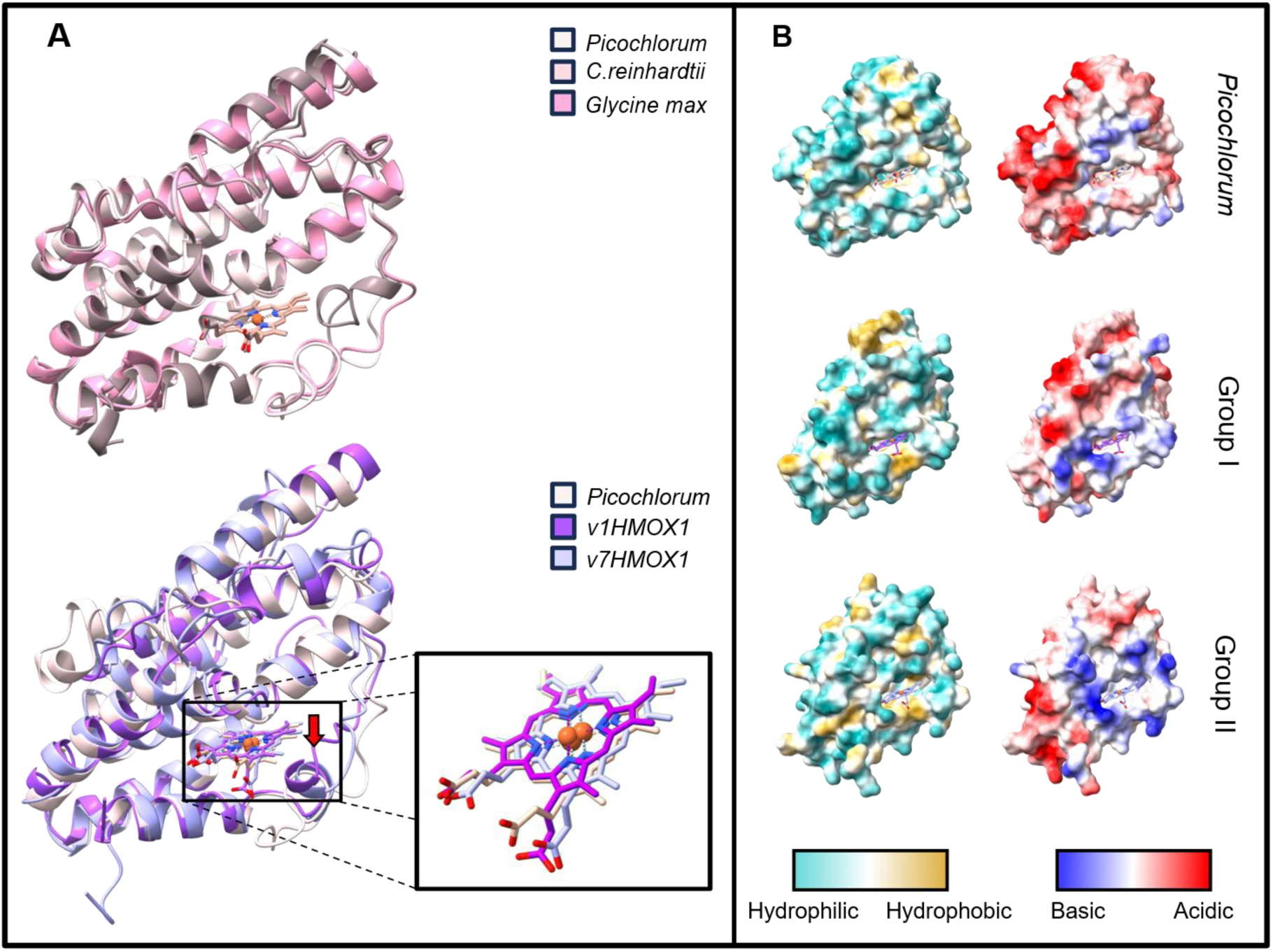
Overall structural features of green lineage and Viral HMOX1 predictions. The top pane of **panel A** shows superimposed structurally predicted HMOX1 proteins from *Chlorophytes Picochlorum sp.* and *Chlamydomonas reinhardtii* superimposed with the experimentally determined structure of *Glycine max.* HMOX1 in complex with heme (PDB ID: 7cka). The bottom pane shows the superimposed predicted host protein from *Picochlorum sp.* with v1HMOX1, a member of Group I viral enzymes and v7HMOX1 of Group II enzymes. The red arrow indicates potential proximal loop restructuration. The insert in this panel is a close up of the orientation of the heme substrate in the different enzymes. **Panel B** shows, from top to bottom, the predicted hydrophobicity and electrostatic potential for the *Picochlorum sp.,* the v1HMOX1 representative for Group I and the v7HMOX1 for Group II proteins.

Like in all HMOX1 structures, the overall architecture is composed of eight α-helices with the heme located in the pocket between the N-terminal and seventh helix of all viral structures (Fig. 3A). Conservation and divergence of structural features are reported in Table S4. The heme iron shows the same coordination as seen by the imidazole group of His30 of GmHMOX1 on the proximal side while the propionate groups are similarly pointing out from the pocket in all enzymes (Fig. 3B, and Table S4A) The conserved catalytic residues and proximal histidine–heme coordination geometry were maintained across Groups I and II, supporting preservation of the canonical catalytic scaffold (Fig. S3 and Table S4A). Despite this overall structural conservation, local differences were observed around the heme-binding environment. On the proximal side, the plant enzymes, GmHMOX1 and AtHMOX1 (*Arabidopsis thaliana*; PDB ID: 7EQH) have been described as having a hole which runs from the a-meso carbon to the heme pocket which is absent in mammalian HMOX1 enzymes. This conserved feature has been suggested to confer the specificity of plant enzymes. Interestingly, the viral enzymes show an even greater opening in this area (Fig. 3B and 3C). Another noticeable feature in plant enzymes is the presence of a set of four 3_10_ helices (residues 45–47, 50–52, 84–89, and 172–174 in GmHMOX1) which seem to have disappeared in vHMOX1 models (Fig. 3A, Fig. S3). On the contrary, the viral enzymes have acquired interesting features in the vicinity of the heme binding site which are absent from structurally determined plant enzymes but also from the predicted *Picochlorum sp.* host. An extended α-helical segment forming a 3_10_ helix, otherwise absent in plant structures and *Chlorophyte* predictions, reshapes the proximal region of the viral enzymes heme pocket (Fig. 2A and Fig. 3A). In addition, both viral groups displayed a more electropositive pocket entrance than PicoHMOX1, whereas hydrophobicity patterns were conserved within each viral group (Fig. 2B). Together, these observations indicate local divergence of the heme-binding environment between vHMOX1. Specific remodeling of the propionate environment is also observed in both viral enzyme groups. The h-bonding network to two conserved Arg residues, Arg74 and Arg237 in *Picochlorum* are conserved but an additional tyrosine residue, Tyr109 (Group I nomenclature) is also involved the coordinating the heme (Fig. S3, Table. S4A) while preserving the overall heme oxygenase type 1 catalytic architecture. The local propionate environment is therefore coordinated by an additional number of H-bonds on one propionate while the other is more loosly coordinate than in host HMOX1.

**Figure 3:**
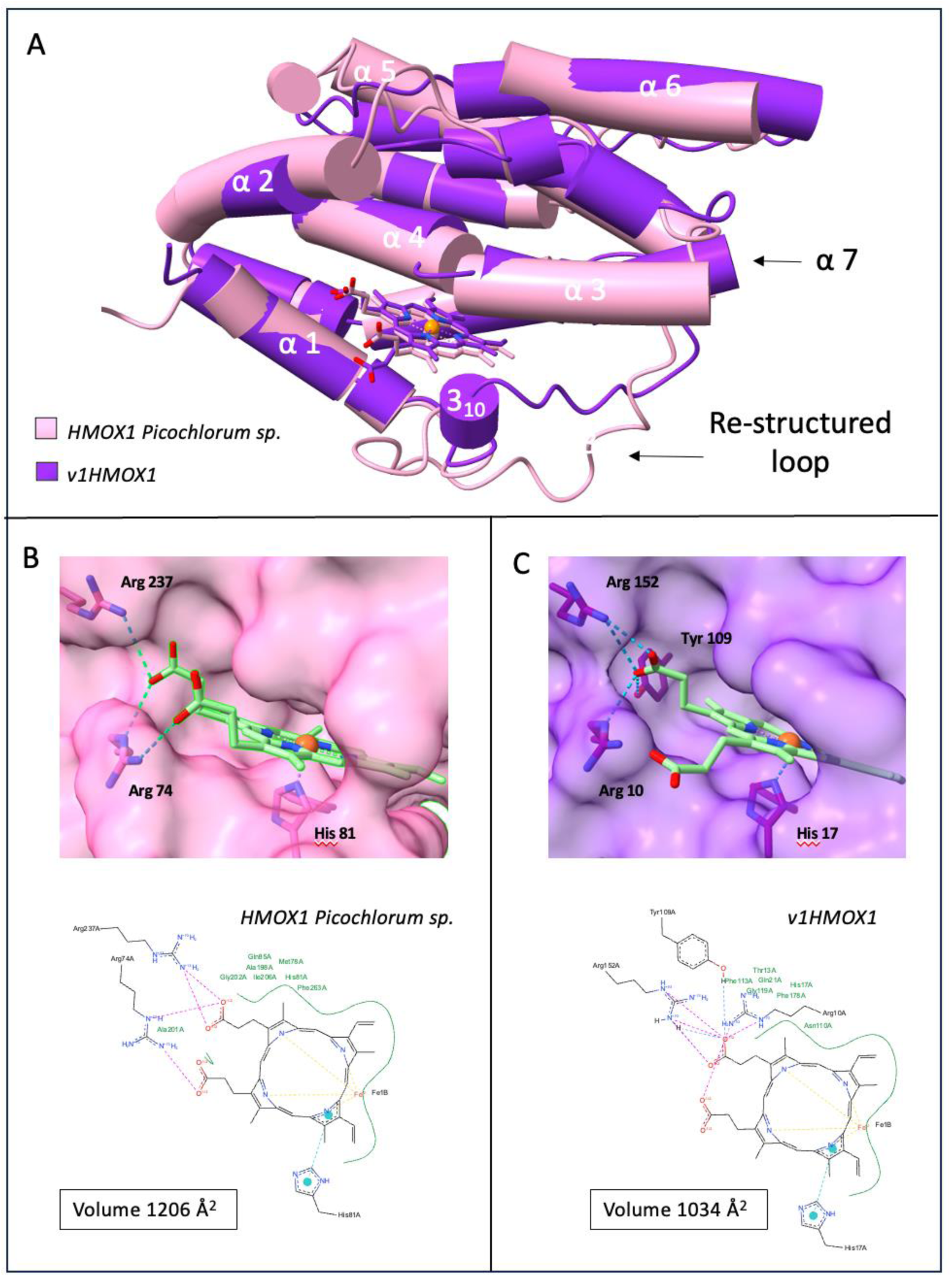
Host and vHMOX1 heme pocket binding analysis. **A:** In HMOX1 enzymes the binding pocket is located at the interface of α1, α2, α3, α4 and α7. The pocket is also delimited by an unstructured loop which shows the appearance of a 310 helix in the predicted viral enzymes which is absent from host enzyme predictions. **B:** The predicted heme binding pocket of *Picochlorum sp.* shows a volume of 1206 Å and a H-bonding network of the propionyl groups established by two Arg residues, Arg237 and Arg74. (**C**) In the case of viral enzymes show a similar H-bonding network of the propionyl groups established by two Arg residues (Arg152 andArg10) but an additional H-bond is present with Tyr109, while the pocket size shows a volume of 1034 Å. For simplicity only v1 is shown.

### Chloroplastic ferredoxin vHMOX1 interaction predictions

In photosynthetic organisms, ferredoxin is the physiological electron donor and is exclusively found in the chloroplast. A particularly important study solved the crystal structure of soybean HMOX1 (GmHMOX1, PDB ID: 7CKA) and mapped the ferredoxin interaction using NMR and ITC (Tohda *et al*. 2021). The ferredoxin-binding interface was identified, but the authors did not report an experimentally determined structure of the GmHMOX1–ferredoxin complex, but rather a model (PDB ID: 9A8X). Along those lines, predicting the feasibility of vHMOX-ferredoxin complexes using the chloroplastic homologue ferrodoxin-3 from *Picochlorum sp.* BPE23 *c*an provide insight into whether or not this protein can be compatible as an electron donor for viral enzymes specifically.

The available structural *Glycine max* data suggest that the two native heme propionates contribute differently in the HMOX1-ferredoxin interaction and catalytic mechanism. One propionate forms a key ionic interaction with a conserved basic residue (Arg23 in GmHMOX1). This interaction is important for fixing the heme orientation (Tohda *et al*. 2021) and can be seen to be conserved and even reinforced by a Tyr interaction in viral enzymes (Fig. 3C). The second propionate also participates in hydrogen-bonding and electrostatic interactions that further stabilize the correct heme orientation and catalytic geometry in *Glycine max* (Tohda *et al*. 2021). Regarding the GmHMOX1–ferredoxin interaction itself, there is currently no evidence that ferredoxin directly recognizes both heme propionates. The propionates are thought to be important mainly because they maintain the correct heme orientation and active-site architecture required for electron transfer and catalysis.

The ferredoxin-binding surface appears to be primarily a protein–protein interaction involving charged residues on the GmHMOX1 surface rather than direct contact with the heme propionates. While the ferredoxin orientation is predicted as different in vHMOX1/ferredoxin structures than in *Glycine max HMOX1/Zea Mays* ferredoxin structure (Fig. 4), the overall surface charge remains compatible in the viral complexes, suggesting ferredoxin as an electron donor for vHMOX1 enzymes. A closer look at the predicted distances between heme (HEM) and the FE2/S2 iron sulfur cluster (FES) indicates a distance between 6 and 8 Å, compatible with electron transfer (Fig. 4B and C). Interestingly, the FES molecule in viral structures of both Group I and Group II enzymes are predicted to be positioned at the same distance relative to each other and orientated towards only one of the two propionates, the loosely coordinated one in both cases (Fig. 3C).

**Figure 4:**
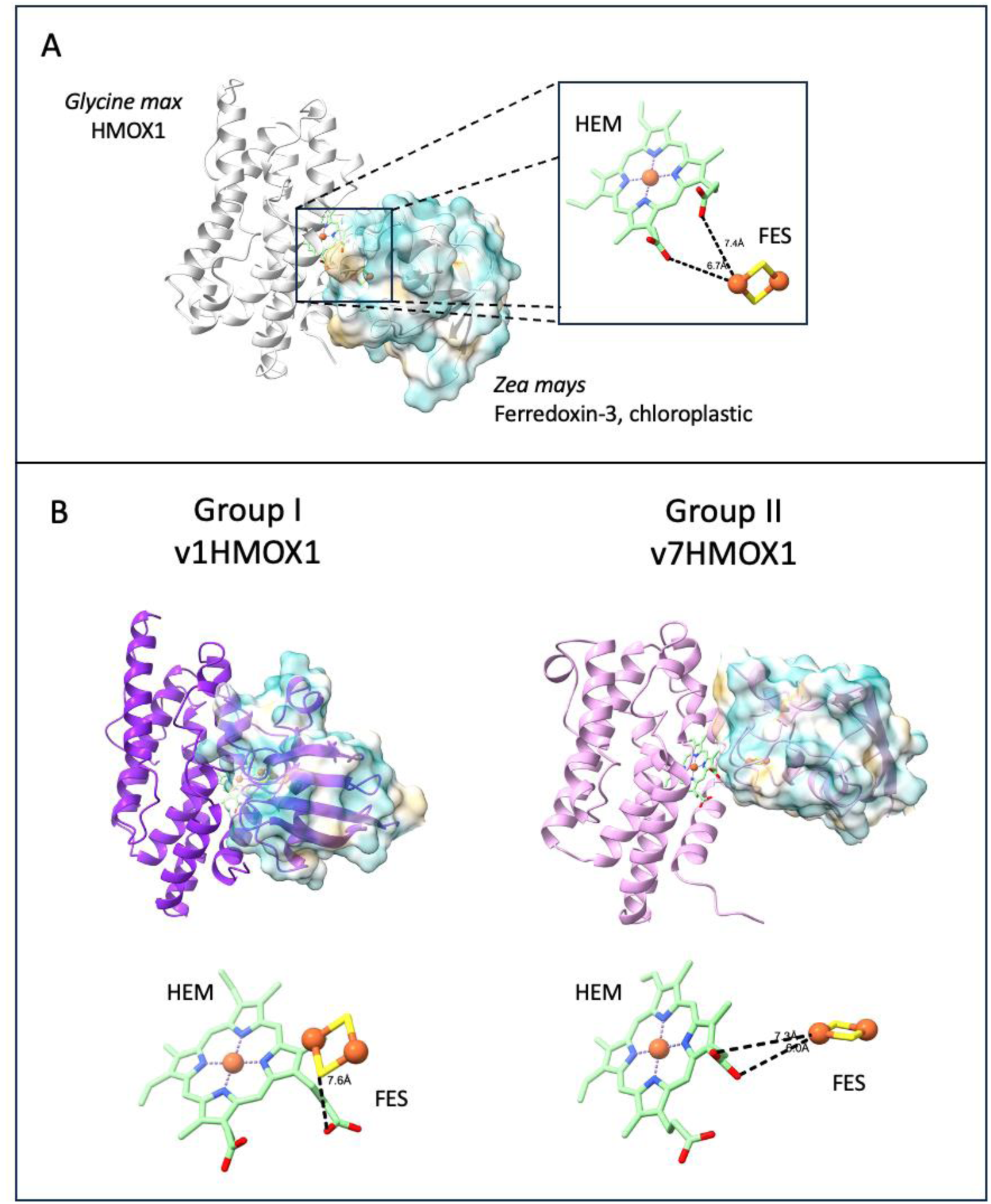
vHMOX1 and ferredoxin binding analysis. **A**: Structural representation of the predicted *Glycine max* HMOX1 and *Zea Mays* ferredoxin interaction based on ITC and NMR interaction studies (PDB ID: 9A8X). The HMOX1 structure is represented as a cartoon while the ferredoxin cartoon structure is encompassed in its hydrophobic surface representation. The heme and iron-sulfur cluster are represented as sticks and the inlet shows a close up including the distances between both ligands. **B** and **C** show the same information for Group I and Group II HMOX1 enzymes, respectively.

### Overall vPcyA structural analysis

Several experimentally determined PcyA structures are present in the Protein Data Bank (https://www.rcsb.org/) mostly of cyanobacterial enzymes. A single *Chlorophyte* PcyA structure from the green algae *Chlamydomonas reinhardtii* has been determined as part of the multi-protein Ycf2-FtsHi chloroplast protein import motor, which the author suggested could be an unknown moonlighting function for PcyA (Liang *et al*. 2024). In all cases PcyA is folded in a three-layer α/β/α sandwich structure. While the *Chlamydomonas reinhardtii* structure is in its Apo form, BV bound structures show a conserved basic patch present on the PcyA surface near the BV molecule entrance suggested to provide a binding site for the acidic ferredoxin electron donor, allowing direct transfer of electrons to BV via its propionate groups (Hagiwara *et al*. 2010; Wada *et al*. 2010). The latter is positioned in a cyclic conformation between the β-sheet and C-terminal α-helices with its two propionate groups extending out towards the basic patch (Fig. 5A, top panel). As shown in Fig. 1B and Fig. S2, algal PcyA structures have long N-terminal extensions which are absent in cyanobacterial and viral enzymes, almost 100 amino acids for *Picochlorum sp*. and approximately 150 amino acids. in *Chlamydomonas reinhardtii*. As previously mentioned, this extension includes the cTP in *Chlorophytes* but with an average length of 30-50 amino acids, the length of these termini are likely involved in other functions, as it is suggested by protein-protein interactions in the Ycf2-FtsHi chloroplast protein import motor (Liang *et al*. 2024).

**Figure 5:**
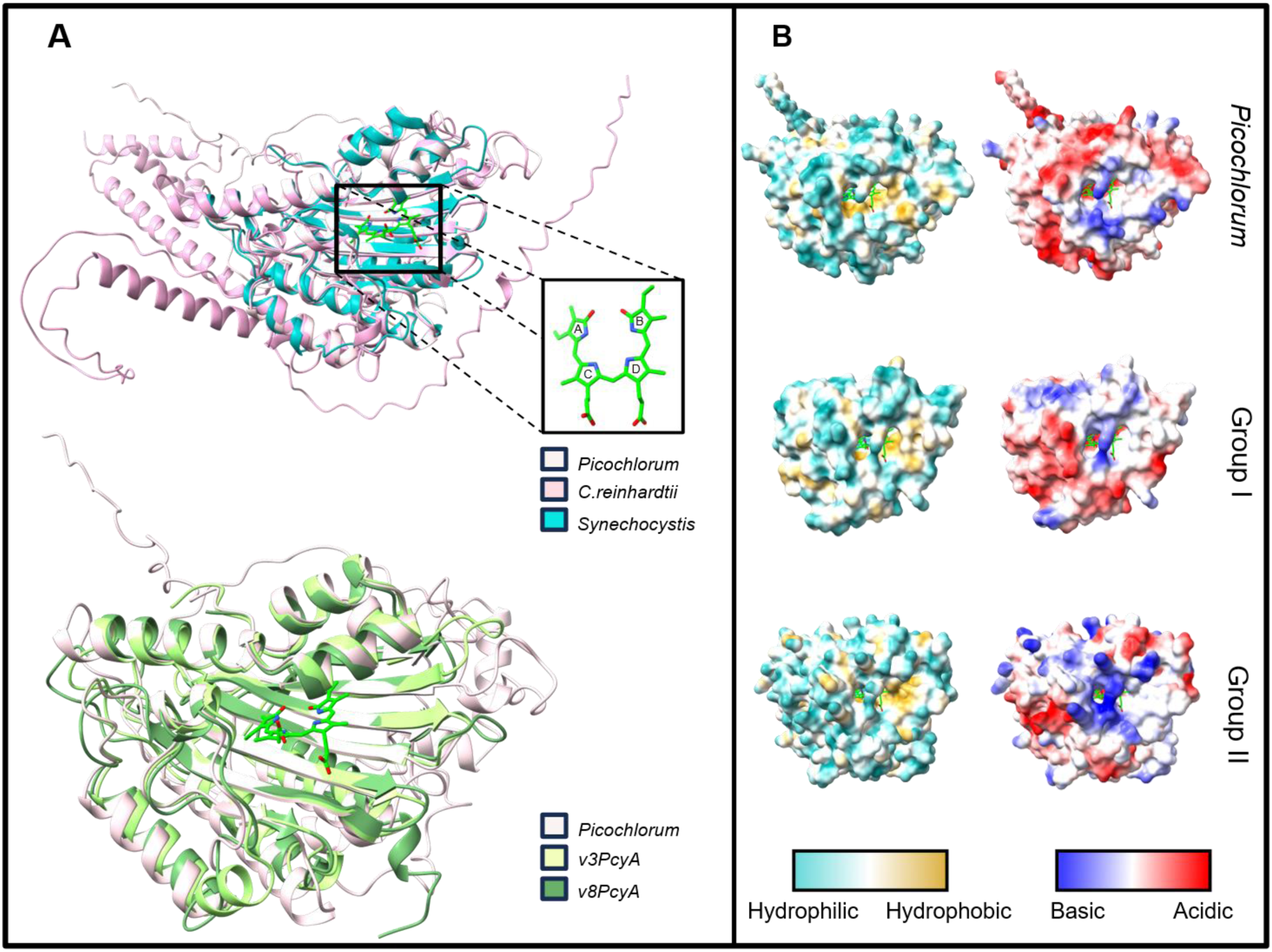
Analysis of Viral PcyA structural predictions. The top pane of **panel A** shows superimposed structurally predicted PcyA proteins from *Chlorophytes Picochlorum sp.* and *Chlamydomonas reinhardtii* superimposed with the experimentally determined structure of *Synechocystis sp.* PcyA which is complexed with biliverdin IXα (PDB ID: 2D1E). The insert in this pane represents the biliverdin IXα substrate with the positioning of the A through D pyrrole rings. The bottom pane shows the superimposed predicted host protein from *Picochlorum sp.* with v3PcyA, a member of Group I viral enzymes and V8PcyA of Group II enzymes, with the BVIX orientation from the SynPcyA structure. **Panel B** shows, from top to bottom, the predicted hydrophobicity and electrostatic potential for the *Picochlorum sp.,* the v3PcyA representative for Group I and the v8PcyA for Group II proteins.

The sequential reduction of the D- and A-rings on the BV is controlled by a central Asp residue located between the two reduction sites that plays its role by changing conformations during the reaction. An extensive hydrophobic contact network has been proposed between PcyA and BV to stabilize its binding with all hydrophilic functional groups of BV being hydrogen-bonded and/or salt-bridged to PcyA (Hagiwara *et al*. 2010). Sequence alignment of viral and algal PcyA shows high conservation of residues involved in biliverdin (BV) binding (Fig. S4). Of the eleven residues reported to participate in substrate binding (Table. S4B), ten were strictly conserved across all viral sequences. The only exception is Val90, which was substituted in variants v7 and v8 (vPCyA Group II members) but remained conserved in all other vPcyA. Overall, the strong conservation of the BV-binding pocket suggests that substrate recognition and binding are preserved in the viral enzymes. The five residues directly involved in catalysis (Table S4B) were present in every viral sequence analysed. Notably, the key proton-donating residues Glu76, Asp105, and Tyr212, which drive the sequential reduction of the D and A rings of biliverdin, showed complete conservation. These results indicate that the catalytic machinery of viral PcyA is highly conserved, supporting the hypothesis that these enzymes retain the ability to catalyse biliverdin reduction despite their viral origin.

While sequence alignment shows the conservation of all essential and catalytic residues in PcyA orthologs and in our viral enzymes, 3D folding prediction of all 9 viral PcyAs was done (Fig. S6). To compare them, we also predict the 3D structure of *Chlamydomonas reinhardtii*, *Picochlorum sp*. homologue, v3PcyA (Group I representative) and v8PcyA (Group II representative) and align them over the experimental PcyA model of *Synechocystis sp*. (PDB ID: 2D1E). The top pane of panel A of Fig. 5A shows the superimposed structures of the *Chlorophytes* exhibiting long N-terminal extensions in the background. These superpositioned models show that both the BV binding pocket and the basic patch at the entrance (Fig. 5B) present in the crystallographically determined *Synechocystis sp.* structures are highly conserved in the *Chlorophyte* models (Fig. 5A top panel). As such, for our analysis on viral enzymes we compared viral models to the predicted host PcyA model (PicoPcyA), (Fig. 5 bottom pane of Panel A and panel B). The overall core structure of v3PcyA, v8pcyA and PicoPcyA are highly conserved with low RMSD values (Table S3B). The viral PcyA models retained a conserved hydrophobic active-site environment, despite a more positively charged electrostatic surface than the PicoPcyA (Fig. 5B). The BV-binding pocket must provide a balanced environment combining hydrophobic regions to accommodate the tetrapyrrole scaffold and polar or charged residues to stabilize its propionate groups and properly orient the substrate for catalysis. The conservation of these features among viral PcyA models strongly suggests that the structural determinants required for chromophore binding have been maintained despite evolutionary divergence. Minor differences observed at peripheral surface regions may reflect viral-specific adaptations, whereas the preservation of the active-site environment supports the hypothesis that vPcyAs retain the capacity to interact with the BV substrate.

## Discussion

Because HMOX1 and PcyA are nucleus encoded proteins that are then translated in the cytoplasm and addressed to the chloroplast in the model green alga *Chlamydomonas reinhardtii* (Merchant *et al*. 2007; Duanmu *et al*. 2013). TargetP 2.0 analysis predicted no transit peptides in either viral HMOX1 and PcyA proteins. Overall, the N-terminal regions of viral proteins were considerably shorter than those of their *Chlorophyte* counterparts. This difference in length likely contributes to the lack of predicted targeting peptides. We note, however, that CTP prediction tools have limited sensitivity and specificity, especially in algae and in highly divergent viral proteins (Almagro Armenteros *et al*. 2019). In particular, chloroplast transit peptides are highly variable in sequence, often poorly conserved, and can range broadly in length (typically ∼30–80 amino acids), which complicates the reliability of *in silico* detection. Therefore, the absence of predicted CTPs does not exclude chloroplast targeting, as localization could rely on atypical, cryptic, or unusually short targeting signals, or alternative mechanisms that remain to be determined. Alternatively, the viral proteins may not be chloroplast-targeted and could instead function in another cellular compartment.

The sequence homology between viral HMOX1 and *Chlorophyte* HMOX1, together with the co- localization of *vhmox1* and *pcya* on viral genomes support the participation in the same metabolic pathways potentially analogous to the chloroplast-localized pathway found in green algae.

Despite the absence of clearly predicted chloroplast transit peptides, chloroplast targeting of viral proteins through alternative trafficking routes has been documented in well-characterized plant virus systems (Gao and Lozano-Durán 2025). For example, the helper component-proteinase of *Potato virus Y* is proposed to reach the chloroplast stroma via endoplasmic-reticulum-derived vesicles followed by protein–protein interactions with the chloroplast targeted Rubisco small subunit, piggybacking its way through the chloroplast envelope (Qin *et al*. 2024). In *Geminiviruses*, C4/AC4 proteins can follow a distinct route involving initial localization at the plasma membrane and subsequent COPI-dependent trafficking to the chloroplast envelope (Tu *et al*. 2015; Medina-Puche *et al*. 2020; Zhao *et al*. 2024). This evidence from other viral systems lends support to chloroplast localisation without dedicated chloroplast target peptides.

Without *in vivo* evidence we cannot exclude that these enzymes operate within the viral factory, a specialized compartment from which *Nucleocytoviricota* (NCLDVs) are thought to reorganize host metabolism. Viral factories are increasingly recognized as dynamic metabolic hubs that actively recruit host proteins, metabolites and organelles required for efficient viral replication (Novoa *et al*. 2005; de Castro *et al*. 2013; Wileman *et al*. 2016). A cytoplasmic localization of vHMOX1 and vPcyA is compatible with potential recruitment to viral replication compartments (viral factories), which are typically derived from host cytoplasm. However, accessibility to the cofactors for the bilin reactions may pose a problem in the viral factory. While NADH/NADPH are likely to be readily accessible through host metabolic processes, the lack of a viral ferredoxin encoded by the viral genome raises the question of how the low-potential electrons required by PcyA could be supplied here. Electron transfer to PcyA, but also to HMOX1, would therefore require an alternative electron-transfer system, potentially recruited from the host. A similar paradox has been noted for the cytosolic HMOX2 of *Chlamydomonas reinhardtii*, whose physiological electron donor remains unknown despite the conservation of heme oxygenase activity (Duanmu *et al*. 2013). The presence of a complete HO1–PcyA pathway further suggests a requirement for a low-potential electron flux, which could be supplied either by host-derived chloroplast ferredoxin or by an as yet unidentified viral or host functional substitute, potentially providing a degree of redox autonomy within the viral factory. Therefore, even if the physiological redox partner(s) of vHMOX1 and vPcyA are not encoded by the viral genome, the establishment of the viral factory could enable the local recruitment and concentration of host-derived cofactors or electron-transfer proteins required to sustain bilin biosynthesis. Supporting this possibility, proteomic analyses of *Mimivirus* viral factories have identified such host proteins involved in redox processes, highlighting the capacity of these compartments to establish specialized metabolic microenvironments (Fridmann-Sirkis *et al*. 2016). In this context, an alternative hypothesis to the viral bilins hijacking host chloroplast metabolism, is a potential role in redox homeostasis or protection against oxidative stress within the viral factory (Brahim Belhaouari *et al*. 2022). Alternatively, our analysis has shown that chloroplastic ferredoxin could be compatible with playing this role in a yet to be determined manner.

Our structural modeling reveals that while viral PcyA enzymes maintain a highly conserved catalytic core and substrate-binding pocket, viral HMOX1 homologs exhibit distinct local active-site remodeling and altered electrostatic surfaces compared to their algal hosts. This indicates that vPcyA has evolved under strict constraints to maintain host-like function, whereas vHMOX1 homologs have diverged, suggesting adaptations for distinct heme-binding dynamics. Such differences could reflect viral modulation of the flux of heme into the bilin-biosynthesis pathway during infection. Possible consequences include competition with host heme-binding processes, altered iron release from heme, or changes in the availability of bilin pigments required for photosynthetic-apparatus biogenesis. These possibilities remain hypotheses that require biochemical and infection-based validation. Despite their conserved catalytic core, viral PcyA proteins are shorter than their algal homologues and lack the N-terminal extensions, including the predicted chloroplast transit peptide and N-terminal regions that may be used for protein-protein interactions and moonlighting functions in algal proteins. The functional consequences of this truncation remain unknown but may reflect adaptation to a different intracellular trafficking route or to the viral infection context. Overall, sequence and structural comparisons support the hypothesis that the viral HMOX1-PcyA module has been retained and may have adapted to support bilin biosynthesis during infection, with vHMOX1 showing greater evolutionary divergence than vPcyA. In our case, AlphaFold has accelerated the screening of these unusual viral genes, pinpointing both structural homologies and differences between AMGs and known host enzymes. In the absence of traditional crystallographic data, high-confidence predictions offer a structural basis to guide downstream biochemical verification of their activity, substrate binding, and intracellular function.

The study undertaken here shows for the first time evidence of a bilin synthesis pathway in viruses belonging to the *Nucleocytoviricota group*, *Phycodnaviridae.* An additional level of analysis, based on *in vivo* approaches, appears to be essential to confirm the role of these AMGs in an infection context. Ideally, determining the expression dynamics of these AMGs throughout the viral cycle and experimentally testing whether their activity alters host physiology would provide direct evidence for their functional contribution to infection Thus, we are currently working on the realization of infections between the two partners to be able to implement such approaches. In the absence of manipulable infectious systems, an alternative is to use the host to overexpress AMGs and analyze their physiological and metabolic effects. Protocols for the genetic transformation of *Picochlorum strains* (*e.g. Picochlorum celeri)* have been developed for the editing of its genome (Krishnan *et al*. 2021, 2023). The adaptation of these protocols to transform the putative *Picochlorum* host of is being developed for the transgenic expression of vHMOX1 and vPcyA. Finally, using a closely related organism, such as *Chlamydomonas reinhardtii,* in which bio-cellular and biomolecular tools are available for transgenic expression, is a relevant alternative also under development for the study of AMGs in an *in vivo* context.

## Acknowledgments

This work was supported by the CNRS EC2CO program (BILIVIR project) and by the Mediterranean Institute for the Environmental Transition (ITEM, TRANSIVIR project; A*MIDEX AMX-19-IET-012). S.Z. was funded through an AMU Inter-ED scholarship.

## References

Abramson J, Adler J, Dunger J, Evans R, Green T, Pritzel A, Ronneberger O, Willmore L, Ballard AJ, Bambrick J, Bodenstein SW, Evans DA, Hung C-C, O’Neill M, Reiman D, Tunyasuvunakool K, Wu Z, Žemgulytė A, Arvaniti E, Beattie C, Bertolli O, Bridgland A, Cherepanov A, Congreve M, Cowen-Rivers AI, Cowie A, Figurnov M, Fuchs FB, Gladman H, Jain R, Khan YA, Low CMR, Perlin K, Potapenko A, Savy P, Singh S, Stecula A, Thillaisundaram A, Tong C, Yakneen S, Zhong ED, Zielinski M, Žídek A, Bapst V, Kohli P, Jaderberg M, Hassabis D, Jumper JM (2024) Accurate structure prediction of biomolecular interactions with AlphaFold 3. Nature 630(8016), 493–500. 10.1038/s41586-024-07487-w

Almagro Armenteros JJ, Salvatore M, Emanuelsson O, Winther O, von Heijne G, Elofsson A, Nielsen H (2019) Detecting sequence signals in targeting peptides using deep learning. Life Science Alliance 2(5), e201900429. 10.26508/lsa.201900429

AltschuP SF, Gish W, Miller W, Myers EW, Lipman DJ (1990) Basic Local Alignment Search Tool. Journal of Molecular Biology 215**, Issue** **3**, 8. 10.1016/S0022-2836(05)80360-2.

Brahim Belhaouari D, Pires De Souza GA, Lamb DC, Kelly SL, Goldstone JV, Stegeman JJ, Colson P, La Scola B, Aherfi S (2022) Metabolic arsenal of giant viruses: Host hijack or self-use? (Eds. RA Fouchier and JW van der Meer) eLife 11, e78674. 10.7554/eLife.78674

de Castro IF, Volonté L, Risco C (2013) Virus factories: biogenesis and structural design. Cellular Microbiology 15(1), 24–34. 10.1111/cmi.12029

Chase EE, Pitot T, Bouchard S, Triplet S, Przybyla C, Gobet A, Desnues C, Blanc G (2024) Viral dynamics in a high-rate algal pond reveals a burst of Phycodnaviridae diversity correlated with episodic algal mortality. (Eds. NK Duggal and FO Aylward) mBio 15(12), e02803–24. 10.1128/mbio.02803-24

Duanmu D, Casero D, Dent RM, Gallaher S, Yang W, Rockwell NC, Martin SS, Pellegrini M, Niyogi KK, Merchant SS, Grossman AR, Lagarias JC (2013) Retrograde bilin signaling enables Chlamydomonas greening and phototrophic survival. Proceedings of the National Academy of Sciences 110(9), 3621–3626. 10.1073/pnas.1222375110

Edgar RC (2004) MUSCLE: multiple sequence alignment with high accuracy and high throughput. Nucleic Acids Research 32(5), 1792–1797. 10.1093/nar/gkh340

Ehrt C, Schulze T, Graef J, Diedrich K, Pletzer-Zelgert J, Rarey M (2025) ProteinsPlus: a publicly available resource for protein structure mining. Nucleic Acids Research 53(W1), W478– W484. https://doi.org/10.1093/nar/gkaf377

Fernández de Castro I, Tenorio R, Risco C (2021) Virus Factories. Encyclopedia of Virology 495–500. 10.1016/B978-0-12-814515-9.00001-1

Fridmann-Sirkis Y, Milrot E, Mutsafi Y, Ben-Dor S, Levin Y, Savidor A, Kartvelishvily E, Minsky A (2016) Efficiency in Complexity: Composition and Dynamic Nature of Mimivirus Replication Factories. Journal of Virology 90(21), 10039–10047. 10.1128/jvi.01319-16

Fuhrman JA (1999) Marine viruses and their biogeochemical and ecological effects. Nature 399(6736), 541–548. 10.1038/21119

Gao M, Lozano-Durán R (2025) Symptom Development in Plant Viral Diseases: What, How, and Why? Annual Review of Phytopathology 63(1), 431–450. 10.1146/annurev-phyto-121323-021434

Hagiwara Y, Sugishima M, Khawn H, Kinoshita H, Inomata K, Shang L, Lagarias JC, Takahashi Y, Fukuyama K (2010) Structural Insights into Vinyl Reduction Regiospecificity of Phycocyanobilin:Ferredoxin Oxidoreductase (PcyA). Journal of Biological Chemistry 285(2), 1000–1007. 10.1074/jbc.M109.055632

Holm L, Laiho A, Törönen P, Salgado M (2023) DALI shines a light on remote homologs: One hundred discoveries. Protein Science : A Publication of the Protein Society 32(1), e4519. 10.1002/pro.4519

Krishnan A, Cano M, Karns DA, Burch TA, Likhogrud M, Aqui M, Bailey S, Verruto J, Lambert W, Kuzminov F, Naghipor M, Wang Y, Ebmeier CC, Weissman JC, Posewitz MC (2023) Simultaneous CAS9 editing of cpSRP43, LHCA6, and LHCA7 in Picochlorum celeri lowers chlorophyll levels and improves biomass productivity. Plant Direct 7(9), e530. 10.1002/pld3.530

Krishnan A, Likhogrud M, Cano M, Edmundson S, Melanson JB, Huesemann M, McGowen J, Weissman JC, Posewitz MC (2021) Picochlorum celeri as a model system for robust outdoor algal growth in seawater. Scientific Reports 11(1), 11649. 10.1038/s41598-021-91106-5

Ledermann B, Béjà O, Frankenberg-Dinkel N (2016) New biosynthetic pathway for pink pigments from uncultured oceanic viruses. Environmental Microbiology 18(12), 4337–4347. 10.1111/1462-2920.13290

Liang K, Zhan X, Li Y, Yang Y, Xie Y, Jin Z, Xu X, Zhang W, Lu Y, Zhang S, Zou Y, Feng S, Wu J, Yan Z (2024) Conservation and specialization of the Ycf2-FtsHi chloroplast protein import motor in green algae. Cell 187(20), 5638–5650.e18. 10.1016/j.cell.2024.08.002

Lindell D, Jaffe JD, Johnson ZI, Church GM, Chisholm SW (2005) Photosynthesis genes in marine viruses yield proteins during host infection. Nature 438(7064), 86–89. 10.1038/nature04111

Lu S, Wang J, Chitsaz F, Derbyshire MK, Geer RC, Gonzales NR, Gwadz M, Hurwitz DI, Marchler GH, Song JS, Thanki N, Yamashita RA, Yang M, Zhang D, Zheng C, Lanczycki CJ, Marchler-Bauer A (2020) CDD/SPARCLE: the conserved domain database in 2020. Nucleic Acids Research 48(D1), D265–D268. 10.1093/nar/gkz991

Marchler-Bauer A, Bryant SH (2004) CD-Search: protein domain annotations on the fly. Nucleic Acids Research 32(Web Server), W327–W331. 10.1093/nar/gkh454

Medina-Puche L, Tan H, Dogra V, Wu M, Rosas-Diaz T, Wang L, Ding X, Zhang D, Fu X, Kim C, Lozano-Duran R (2020) A Defense Pathway Linking Plasma Membrane and Chloroplasts and Co-opted by Pathogens. Cell 182(5), 1109–1124.e25. 10.1016/j.cell.2020.07.020

Merchant SS, Prochnik SE, Vallon O, Harris EH, Karpowicz SJ, Witman GB, Terry A, Salamov A, Fritz-Laylin LK, Maréchal-Drouard L, Marshall WF, Qu L-H, Nelson DR, Sanderfoot AA, Spalding MH, Kapitonov VV, Ren Q, Ferris P, Lindquist E, Shapiro H, Lucas SM, Grimwood J, Schmutz J, Cardol P, Cerutti H, Chanfreau G, Chen C-L, Cognat V, Croft MT, Dent R, Dutcher S, Fernández E, Ferris P, Fukuzawa H, González-Ballester D, González-Halphen D, Hallmann A, Hanikenne M, Hippler M, Inwood W, Jabbari K, Kalanon M, Kuras R, Lefebvre PA, Lemaire SD, Lobanov AV, Lohr M, Manuell A, Meier I, Mets L, Mittag M, Mittelmeier T, Moroney JV, Moseley J, Napoli C, Nedelcu AM, Niyogi K, Novoselov SV, Paulsen IT, Pazour G, Purton S, Ral J-P, Riaño-Pachón DM, Riekhof W, Rymarquis L, Schroda M, Stern D, Umen J, Willows R, Wilson N, Zimmer SL, Allmer J, Balk J, Bisova K, Chen C-J, Elias M, Gendler K, Hauser C, Lamb MR, Ledford H, Long JC, Minagawa J, Page MD, Pan J, Pootakham W, Roje S, Rose A, Stahlberg E, Terauchi AM, Yang P, Ball S, Bowler C, Dieckmann CL, Gladyshev VN, Green P, Jorgensen R, Mayfield S, Mueller-Roeber B, Rajamani S, Sayre RT, Brokstein P, Dubchak I, Goodstein D, Hornick L, Huang YW, Jhaveri J, Luo Y, Martínez D, Ngau WCA, Otillar B, Poliakov A, Porter A, Szajkowski L, Werner G, Zhou K, Grigoriev IV, Rokhsar DS, Grossman AR (2007) The Chlamydomonas Genome Reveals the Evolution of Key Animal and Plant Functions. *Science (New York*, N.Y.) 318(5848), 245–250. 10.1126/science.1143609

Netherton C, Moffat K, Brooks E, Wileman T (2007) A Guide to Viral Inclusions, Membrane Rearrangements, Factories, and Viroplasm Produced During Virus Replication. Advances in Virus Research 70, 101–182. 10.1016/S0065-3527(07)70004-0

Netherton CL, Wileman T (2011) Virus factories, double membrane vesicles and viroplasm generated in animal cells. Current Opinion in Virology 1(5), 381–387. 10.1016/j.coviro.2011.09.008

Novoa RR, Calderita G, Arranz R, Fontana J, Granzow H, Risco C (2005) Virus factories: associations of cell organelles for viral replication and morphogenesis. Biology of the Cell 97(2), 147–172. 10.1042/BC20040058

Pettersen EF, Goddard TD, Huang CC, Meng EC, Couch GS, Croll TI, Morris JH, Ferrin TE (2021) UCSF ChimeraX: Structure visualization for researchers, educators, and developers. Protein Science 30(1), 70–82. 10.1002/pro.3943

Qin L, Liu H, Liu P, Jiang L, Cheng X, Li F, Shen W, Qiu W, Dai Z, Cui H (2024) Rubisco small subunit (RbCS) is co-opted by potyvirids as the scaffold protein in assembling a complex for viral intercellular movement. PLOS Pathogens 20(3), e1012064. 10.1371/journal.ppat.1012064

Rosenwasser S, Ziv C, Creveld SG van, Vardi A (2016) Virocell Metabolism: Metabolic Innovations During Host-Virus Interactions in the Ocean. Trends in Microbiology 24(10), 821–832. 10.1016/j.tim.2016.06.006

Suttle CA (2005) Viruses in the sea. Nature 437(7057), 356–361. 10.1038/nature04160

Suttle CA (2007) Marine viruses--major players in the global ecosystem. Nature Reviews. Microbiology 5(10), 801–812. 10.1038/nrmicro1750

Tardif M, Atteia A, Specht M, Cogne G, Rolland N, Brugière S, Hippler M, Ferro M, Bruley C, Peltier G, Vallon O, Cournac L (2012) PredAlgo: A New Subcellular Localization Prediction Tool Dedicated to Green Algae. Molecular Biology and Evolution 29(12), 3625–3639. 10.1093/molbev/mss178

Thompson LR, Zeng Q, Kelly L, Huang KH, Singer AU, Stubbe J, Chisholm SW (2011) Phage auxiliary metabolic genes and the redirection of cyanobacterial host carbon metabolism. Proceedings of the National Academy of Sciences 108(39). 10.1073/pnas.1102164108

Tohda R, Tanaka H, Mutoh R, Zhang X, Lee Y-H, Konuma T, Ikegami T, Migita CT, Kurisu G (2021) Crystal structure of higher plant heme oxygenase-1 and its mechanism of interaction with ferredoxin. Journal of Biological Chemistry 296, 100217. 10.1074/jbc.RA120.016271

Tu Y, Jin Y, Ma D, Li H, Zhang Z, Dong J, Wang T (2015) Interaction between PVY HC-Pro and the NtCF1β-subunit reduces the amount of chloroplast ATP synthase in virus-infected tobacco. Scientific Reports 5(1), 15605. 10.1038/srep15605

Wada K, Hagiwara Y, Yutani Y, Fukuyama K (2010) One residue substitution in PcyA leads to unexpected changes in tetrapyrrole substrate binding. Biochemical and Biophysical Research Communications 402(2), 373–377. 10.1016/j.bbrc.2010.10.037

Wang Y, Hou C, Rockwell NC, Brzezowski P, Zhang W, Huang X, Fan Q, Wang Y, Grimm B, Lagarias JC, Duanmu D (2026) Covalent phytobilin adducts of GUN4 implicate a photoprotective mechanism in chlorophyll biosynthesis. Proceedings of the National Academy of Sciences 123(28), e2533100123. 10.1073/pnas.2533100123

Waterhouse A, Bertoni M, Bienert S, Studer G, Tauriello G, Gumienny R, Heer FT, de Beer TAP, Rempfer C, Bordoli L, Lepore R, Schwede T (2018) SWISS-MODEL: homology modelling of protein structures and complexes. Nucleic Acids Research 46(W1), W296–W303. 10.1093/nar/gky427

Waterhouse AM, Procter JB, Martin DMA, Clamp M, Barton GJ (2009) Jalview Version 2—a multiple sequence alignment editor and analysis workbench. Bioinformatics 25(9), 1189– 1191. 10.1093/bioinformatics/btp033

Wegner H, Roitman S, Kupczok A, Braun V, Woodhouse JN, Grossart H-P, Zehner S, Béjà O, Frankenberg-Dinkel N (2024) Identification of Shemin pathway genes for tetrapyrrole biosynthesis in bacteriophage sequences from aquatic environments. Nature Communications 15(1), 8783. 10.1038/s41467-024-52726-3

Wileman T, Netherton CL, Powell PP (2016) Virus Factories and Mini-Organelles Generated for Virus Replication. Encyclopedia of Cell Biology 819–827. 10.1016/B978-0-12-394447-4.20102-8

Wittkopp TM, Schmollinger S, Saroussi S, Hu W, Zhang W, Fan Q, Gallaher SD, Leonard MT, Soubeyrand E, Basset GJ, Merchant SS, Grossman AR, Duanmu D, Lagarias JC (2017) Bilin-Dependent Photoacclimation in Chlamydomonas reinhardtii. The Plant Cell 29(11), 2711–2726. 10.1105/tpc.17.00149

Zhang W, Willows RD, Deng R, Li Z, Li M, Wang Y, Guo Y, Shi W, Fan Q, Martin SS, Rockwell NC, Lagarias JC, Duanmu D (2021) Bilin-dependent regulation of chlorophyll biosynthesis by GUN4. Proceedings of the National Academy of Sciences 118(20), e2104443118. 10.1073/pnas.2104443118

Zhao W, Ji Y, Zhou Y, Wang X (2024) Geminivirus C4/AC4 proteins hijack cellular COAT PROTEIN COMPLEX I for chloroplast targeting and viral infections. Plant Physiology 196(3), 1826–1839. 10.1093/plphys/kiae436

