## Supplemental Figures and Tables for "Analysis of *Phycodnaviridae* Tetrapyrrole Enzymes: Subcellular Localization and Functional Divergence from Putative Host Homologs"

### Supplementary figures and tables for : “Analysis of *Phycodnaviridae* Tetrapyrrole Enzymes: Subcellular Localization and Functional Divergence from Putative Host Homologs”

Authors : Steven Zehnacker, Stefano Caffarri, Guillaume Blanc, Xenie Johnson and Marina I. Siponen

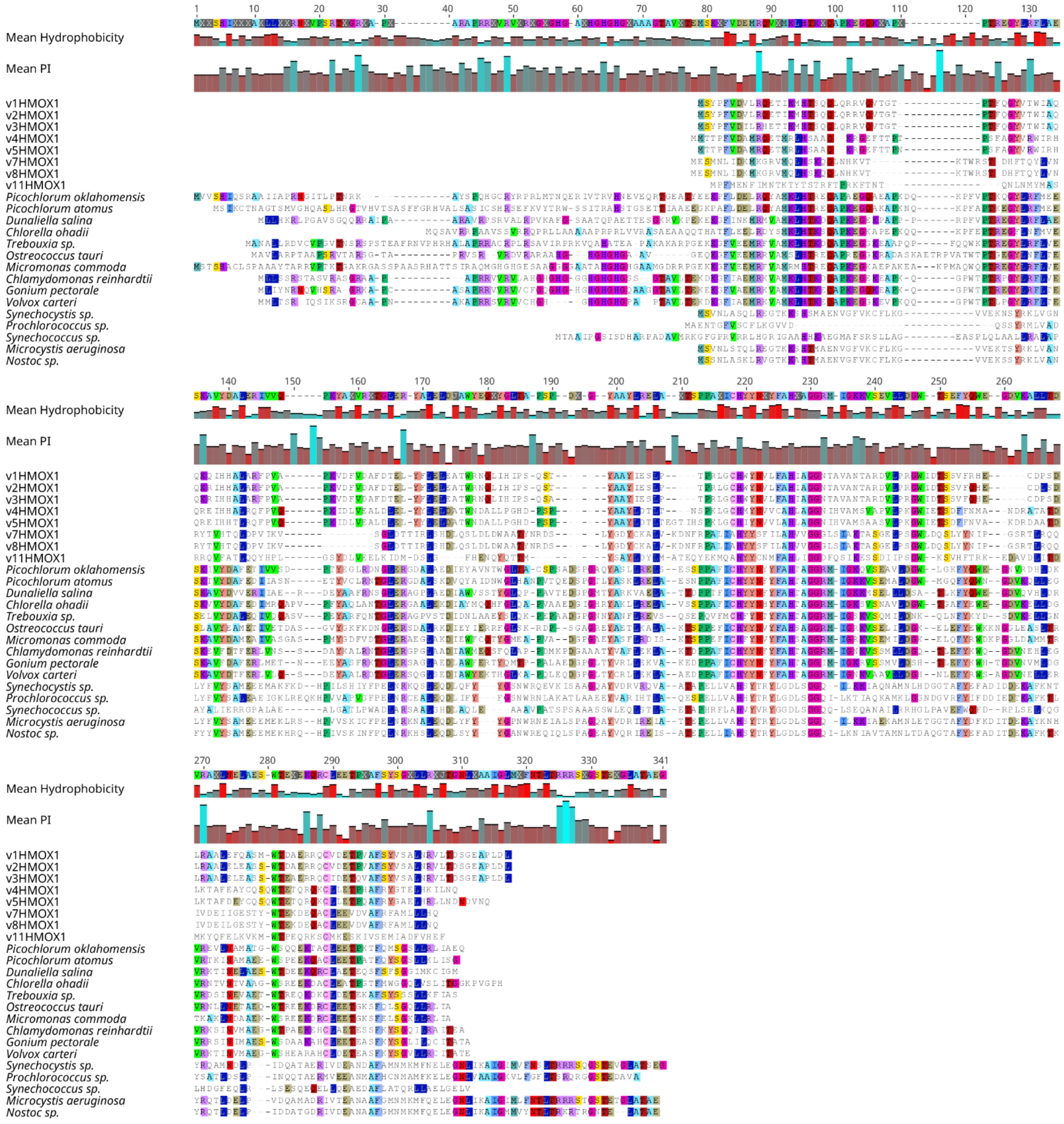

**Figure S1 : Multiple sequence alignment of HMOX1 homologs.** Multiple alignment containing protein sequences from viral, algal, and cyanobacterial HMOX1 homologs. Sequence identifiers and accession numbers are provided in Supplementary Table S1A.

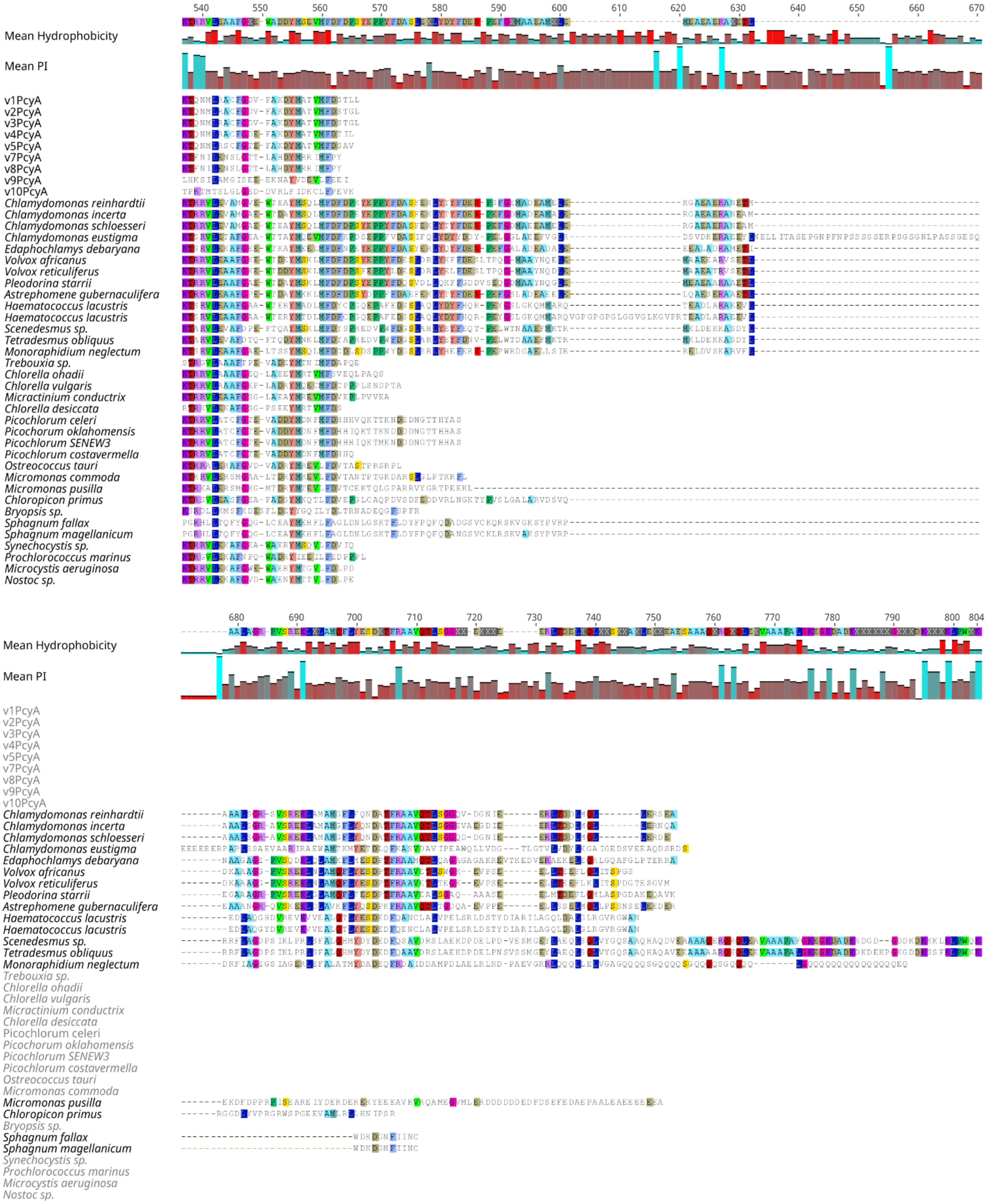


**Figure S2 : Multiple sequence alignment of PcyA homologs.** Multiple alignment containing protein sequences from viral, algal, and cyanobacterial PcyA homologs. Sequence identifiers and accession numbers are provided in Supplementary Table S1B.

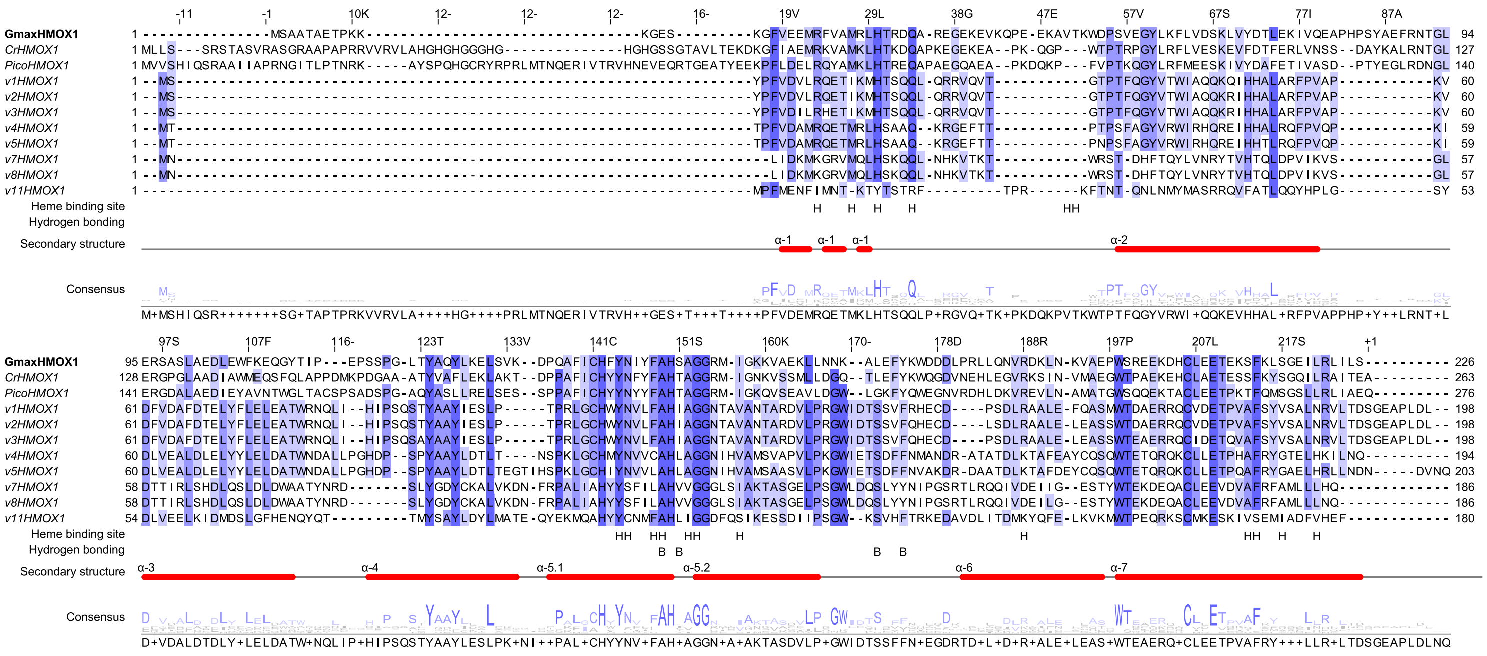

**Figure S3 Sequence alignment of HMOX1 viral enzymes :** The reference sequence used to annotate viral sequences is that of *Glycine max* (GmaxHMOX1). The sequences of the "reference" enzymes correspond to those of *C. reinhardtii* (CrHMOX1) and *Picochlorum*, the presumed host of the viruses. Secondary structural elements are annotated in red (alpha helices). Catalytic residues of GmaxHMOX1 are denoted by an H for heme binding sites and residues involved in hydrogen bridges are indicated by a B.

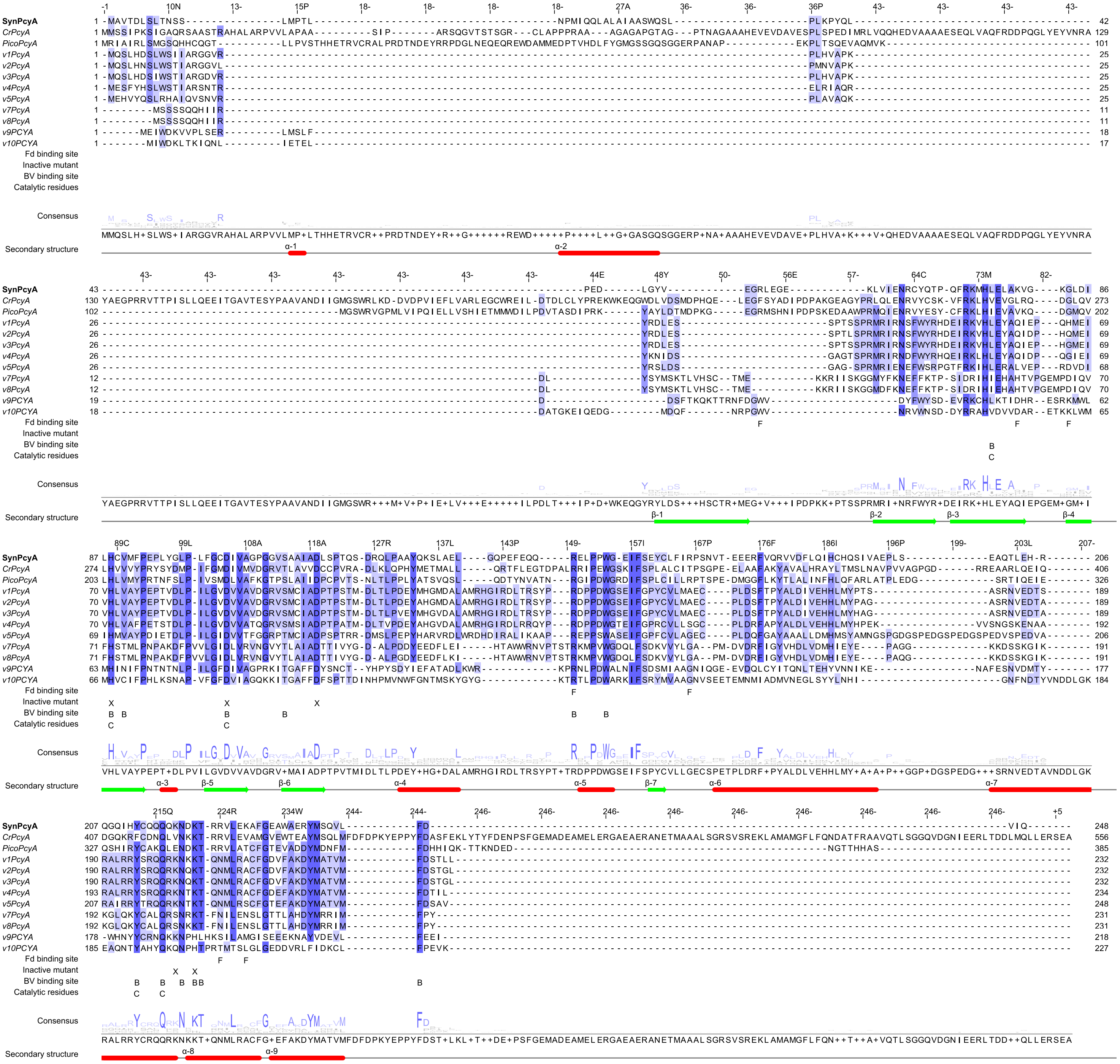


**Figure S4: Sequence alignment of PcyA viral enzymes :** The reference sequence used to annotate viral sequences is that of *Synechocystis* (SynPcyA). The sequences of the "reference" enzymes correspond to those of *C. reinhardtii* (CrPcyA) and *Picochlorum* (PicoPcyA), the presumed host of the viruses. Secondary structural elements are annotated in red (alpha helices) and green (beta sheet). Biliverdin binding sites are denoted by a B, catalytic residues of SynPcyA by a C and mutations that inactivate enzyme function by an X.

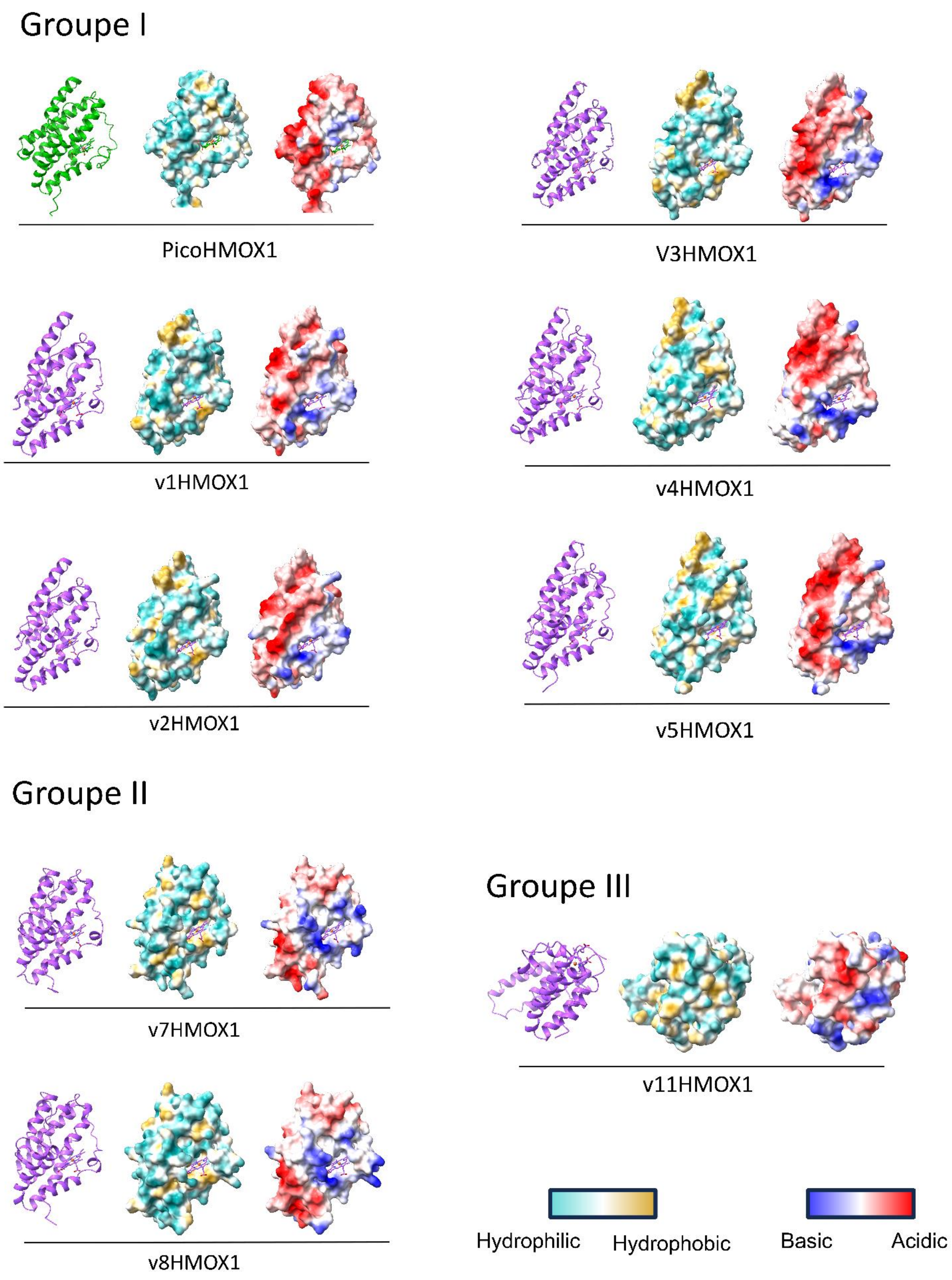
Groupe I
PicoHMOX1
V3HMOX1
v1HMOX1
v4HMOX1
v2HMOX1
v5HMOX1
Groupe II
v7HMOX1
v8HMOX1
Groupe III
v11HMOX1
Hydrophilic
Hydrophobic
Basic
Acidic

**Figure S5: Structural predictions of viral HMOX1 homologs.** Predicted three-dimensional structures of the viral HMOX1 homologs generated using AlphaFold3 with the heme cofactor included during structure prediction. The left representation shows ribbon representations of each predicted protein with the heme molecule displayed as sticks. The middle panel shows the molecular surfaces colored according to hydrophobicity. The right representation shows the corresponding molecular surfaces colored according to electrostatic potential (red, negative; blue, positive). Based on structural predictions and sequence conservation, the viral proteins segregate into three groups: Group I (v1–v5), Group II (v7–v8), and the highly divergent Group III (v11). While Groups I and II retain the canonical HMOX fold, they exhibit distinct electrostatic and hydrophobic surface properties, whereas Group III displays extensive structural divergence, including a markedly different predicted organization of the heme-binding pocket.

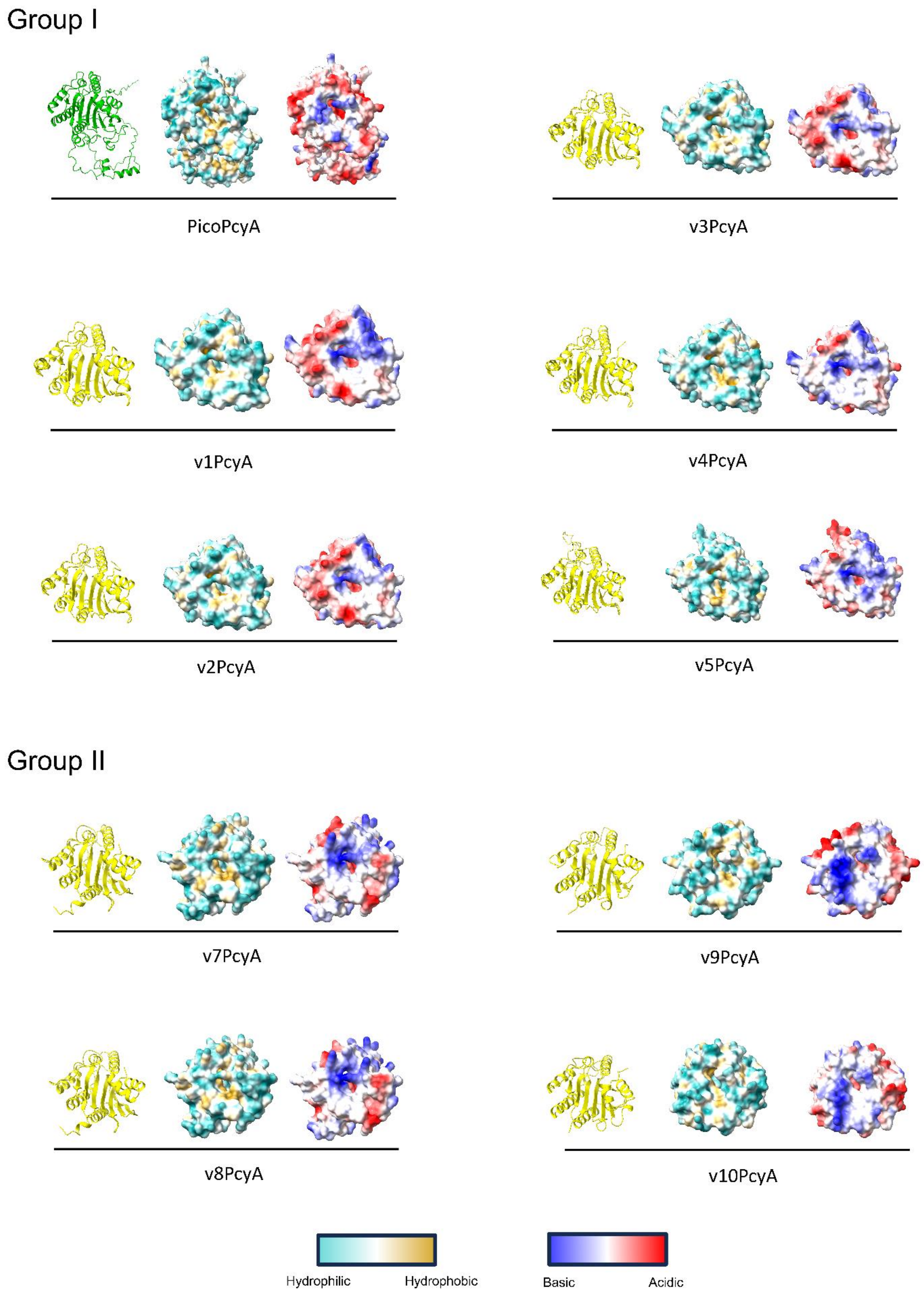


**Figure S6: Structural predictions of viral PcyA homologs.** Predicted three-dimensional structures of the viral PcyA homologs generated using AlphaFold3 with biliverdin IXα included during structure prediction. The left representation shows ribbon representations of each predicted protein. The middle representation shows the corresponding molecular surfaces colored according to electrostatic potential (red, negative; blue, positive). The right panel shows the molecular surfaces colored according to hydrophobicity. Based on structural predictions and sequence conservation, the viral proteins segregate into two groups: Group I (v1–v5) and Group II (v7–v9).

**Table S1A. HMOX1 homolog sequences :** List of HMOX1 homolog sequences used in the analysis, including sequence identifiers/accession numbers and database of origin.

| **Species / sequence** | **Sequence ID / accession** | **Origin** |
|---|---|---|
| *Picochlorum oklahomensis* | Derived from genome assembly downloaded from 10.1093/molbev/msy167 | Foflonker *et al.* 2018 |
| *Picochlorum atomus* | OneKP project database; ID=scaffold-JQFK-2002893-Picochlorum_atomus | OneKP |
| *Dunaliella salina* | KAF5828604.1 | GenBank |
| *Chlorella ohadii* | KAI7836606.1 | GenBank |
| *Trebouxia* sp. A1-2 | KAA6428757.1 | GenBank |
| *Ostreococcus tauri* | OUS45100.1 | GenBank |
| *Micromonas commoda* | XP_002501156.1 | GenBank |
| *Chlamydomonas reinhardtii* | XP_042919914.1 | GenBank |
| *Gonium pectorale* | KXZ53880.1 | GenBank |
| *Volvox carteri* | XP_002951957.1 | GenBank |
| *Synechocystis* sp. PCC 6803 | P72849 | UniProt |
| *Prochlorococcus* sp. | KZR66847.1 | GenBank |
| *Synechococcus* sp. | A0A7T1HP91 | UniProt |
| *Microcystis aeruginosa* | WP_002737675.1 | GenBank |
| *Nostoc* sp. | WP_067775078.1 | GenBank |

**Table S1B. PcyA homolog sequences :** List of PcyA homolog sequences used in the analysis, including sequence identifiers/accession numbers and database of origin.

| Species | Sequence ID / accession | Origin |
|---|---|---|
| *Chlamydomonas reinhardtii* | XP_001693622 | GenBank |
| *Chlamydomonas incerta* | KAG2443466 | GenBank |
| *Chlamydomonas schloesseri* | KAG2443214 | GenBank |
| *Volvox africanus* | GIL51368 | GenBank |
| *Volvox reticuliferus* | GIL75456 | GenBank |
| *Pleodorina starrii* | GLC43951 | GenBank |
| *Astrephomene gubernaculifera* | GFR52683 | GenBank |
| *Edaphochlamys debaryana* | KAG2497231 | GenBank |
| *Chlamydomonas eustigma* | GAX73683 | GenBank |
| *Haematococcus lacustris* | KAJ9522776 | GenBank |
| *Haematococcus lacustris* | KAJ9522471 | GenBank |
| *Scenedesmus* sp. | KAF6265515 | GenBank |
| *Tetradesmus obliquus* | WIA22780 | GenBank |
| *Monoraphidium neglectum* | XP_013902783 | GenBank |
| *Trebouxia* sp. | KAA6427473 | GenBank |
| *Chlorella ohadii* | KAI7844580 | GenBank |
| *Micractinium conductrix* | PSC74514 | GenBank |
| *Chlorella vulgaris* | KAI3425822 | GenBank |
| *Chlorella desiccata* | KAG7668407 | GenBank |
| *Picochlorum celeri* | Derived from Genbank genome assembly GCA_010909725.1 | GenBank |
| *Picochlorum oklahomensis* | Derived from genome assembly downloaded from 10.1093/molbev/msy167 | Foflonker *et al.* 2018 |
| *Picochlorum sp.* SENEW3 | Derived from Genbank genome assembly GCA_011316045.1 | GenBank |
| *Picochlorum costavermella* | Derived from Genbank genome assembly GCA_011316045.1 | GenBank |
| *Ostreococcus tauri* | XP_003078990 | GenBank |
| *Micromonas commoda* | XP_002502590 | GenBank |

| | | |
|---|---|---|
| *Micromonas pusilla* | XP_003059231 | GenBank |
| *Bryopsis* sp. | GMH37566 | GenBank |
| *Sphagnum fallax* | KAH8940247 | GenBank |
| *Sphagnum magellanicum* | KAH9539902 | GenBank |
| *Synechocystis* sp. | Q55891 | UniProt |
| *Synechococcus elongatus* | Q5N5A3 | UniProt |
| *Prochlorococcus marinus* | Q7V7Y8 | UniProt |
| *Microcystis aeruginosa* | B0JPG5 | UniProt |
| *Nostoc* sp. | Q93TN0 | UniProt |

**<u>Table S2A: CTP prediction in HMOX1 sequences :</u>** Prediction of transit peptides using TargetP2.0. Predictions were performed on HMOX1 sequences from putative *Picochlorovirus*, green algae and cyanobacteria. The table summarizes the predicted subcellular localization, along with the corresponding confidence scores for each sequence (cTP : chloroplast transit peptide ; OTHER : No transit peptide detected).

| # ID | Prediction | OTHER | SP | mTP | cTP | lTP | CS Position |
|---|---|---|---|---|---|---|---|
| v1HMOX1 | OTHER | 0.996284 | 0.000335 | 0.002959 | 0.000403 | 0.000018 | |
| v2HMOX1 | OTHER | 0.996071 | 0.000228 | 0.003053 | 0.000617 | 0.000031 | |
| v3HMOX1 | OTHER | 0.995455 | 0.000245 | 0.003542 | 0.000723 | 0.000036 | |
| v4HMOX1 | OTHER | 0.980408 | 0.012775 | 0.005078 | 0.001705 | 0.000033 | |
| v5HMOX1 | OTHER | 0.992847 | 0.005659 | 0.001328 | 0.000162 | 0.000005 | |
| v7HMOX1 | OTHER | 0.990767 | 0.000695 | 0.00836 | 0.000155 | 0.000023 | |
| v8HMOX1 | OTHER | 0.990858 | 0.000711 | 0.008252 | 0.000156 | 0.000023 | |
| v11HMOX1 | OTHER | 0.905816 | 0.000087 | 0.047131 | 0.045711 | 0.001255 | |
| *Chlamydomonas_reinhardtii_A0A2K3D9D4* | cTP | 0.106244 | 0.000293 | 0.407578 | 0.485684 | 0.000201 | CS pos: 28-29. RVL-AH. Pr: 0.1606 |
| *Dunaliella_salina_A0ABQ7G1X4* | cTP | 0.007708 | 0.000002 | 0.489925 | 0.502353 | 0.000012 | CS pos: 37-38. VKA-FG. Pr: 0.4210 |
| *Chlorella_ohadii_A0AAD5DGW3* | cTP | 0.002511 | 0.000067 | 0.038172 | 0.959153 | 0.000097 | CS pos: 35-36. VRA-SA. Pr: 0.8591 |
| *Picochlorum sp. BPE23_KAI8104144.1* | OTHER | 0.524454 | 0.00067 | 0.199233 | 0.274145 | 0.001499 | |
| *Trebouxia_sp._KAA6428757* | cTP | 0.097216 | 0 | 0.014081 | 0.888557 | 0.000145 | CS pos: 56-57. QAH-AT. Pr: 0.3126 |
| *Micromonas_commoda_C1E2W1* | cTP | 0.062917 | 0.000003 | 0.000154 | 0.935786 | 0.00114 | CS pos: 44-45. IRA-QM. Pr: 0.4969 |
| *Ostreococcus_tauri_OUS45100* | cTP | 0.093243 | 0.000024 | 0.018109 | 0.887118 | 0.001506 | CS pos: 34-35. ARA-AH. Pr: 0.2320 |
| *Synechocystis_sp._PCC6803* | OTHER | 0.993564 | 0.00042 | 0.005943 | 0.000058 | 0.000015 | |
| *Prochlorococcus_sp_P72849* | OTHER | 0.999672 | 0.00031 | 0.000017 | 0.000001 | 0.000001 | |
| *Synechococcus_sp.__QPN59229* | OTHER | 0.737133 | 0.00003 | 0.001822 | 0.119707 | 0.141309 | |
| *Microcystis_aeruginosa_A0AB33BP28* | OTHER | 0.996961 | 0.000298 | 0.002656 | 0.000049 | 0.000037 | |
| *Nostoc_sp_BAT56487* | OTHER | 0.967435 | 0.000098 | 0.0309 | 0.001413 | 0.000154 | |

**Table S2B: CTP prediction in PcyA sequences :** Prediction of transit peptides using TargetP2.0. Predictions were performed on PcyA sequences from putative *Picochlorovirus*, green algae and cyanobacteria. The table summarizes the predicted subcellular localization, along with the corresponding confidence scores for each sequence (cTP : chloroplast transit peptide ; mTP : mitochondrial transit peptide ; SP : secretory pathway signal peptide ; OTHER : No transit peptide detected).

| # ID | Prediction | OTHER | SP | mTP | cTP | lTP | CS Position |
|---|---|---|---|---|---|---|---|
| v1PcyA | OTHER | 0.966235 | 0.002172 | 0.024945 | 0.006278 | 0.00037 | |
| v2PcyA | OTHER | 0.878421 | 0.003926 | 0.063774 | 0.052356 | 0.001523 | |
| v3PcyA | OTHER | 0.997722 | 0.000549 | 0.001214 | 0.000459 | 0.000056 | |
| v4PcyA | OTHER | 0.984973 | 0.000242 | 0.012833 | 0.000276 | 0.001677 | |
| v5PcyA | OTHER | 0.916633 | 0.00076 | 0.081407 | 0.00109 | 0.000109 | |
| v7PcyA | OTHER | 0.992829 | 0.000792 | 0.004436 | 0.001824 | 0.000119 | |
| v8PcyA | OTHER | 0.993683 | 0.000771 | 0.004 | 0.001463 | 0.000083 | |
| v9PCYA | OTHER | 0.999619 | 0.00037 | 0.00001 | 0 | 0 | |
| v10PCYA | OTHER | 0.99971 | 0.000204 | 0.000085 | 0 | 0 | |
| *Chlamydomonas_reinhardtii_ XP_001693622* | cTP | 0.050389 | 0.000014 | 0.014313 | 0.935239 | 0.000046 | CS pos: 56-57. PRA-AA. Pr: 0.3204 |
| *Volvox_africanus__GIL51368* | cTP | 0.187859 | 0.000163 | 0.400497 | 0.41145 | 0.000032 | CS pos: 59-60. VRA-ST. Pr: 0.5668 |
| *Chlorella_ohadii__KAI7844580* | OTHER | 0.999856 | 0.000134 | 0.000003 | 0.000006 | 0.000001 | |
| *Picochlorum sp. BPE23_KAI8100569.1* | mTP | 0.404729 | 0.001792 | 0.591103 | 0.002316 | 0.000059 | CS pos: 32-33. VCR-AL. Pr: 0.2713 |
| *Trebouxia_sp._KAA6427473* | cTP | 0.40417 | 0.000027 | 0.023771 | 0.571197 | 0.000835 | CS pos: 73-74. ASQ-TD. Pr: 0.2165 |
| *Micromonas_commoda_XP_002502590* | cTP | 0.149496 | 0.000019 | 0.014944 | 0.835342 | 0.000199 | CS pos: 32-33. ARA-AA. Pr: 0.2938 |
| *Ostreococcus_tauri_XP_003078990* | OTHER | 0.968012 | 0.002592 | 0.000458 | 0.028826 | 0.000112 | |
| *Synechocystis_sp._Q55891* | OTHER | 0.870215 | 0.017623 | 0.001577 | 0.109803 | 0.000782 | |
| *Prochlorococcus_marinus_Q7V7Y8* | SP | 0.246905 | 0.737864 | 0.004823 | 0.009085 | 0.001323 | CS pos: 26-27. STS-GP. Pr: 0.6382 |
| *Synechococcus_elongatus_Q5N5A3* | OTHER | 0.996431 | 0.00067 | 0.002886 | 0.00001 | 0.000002 | |
| *Microcystis_aeruginosa_B0JPG5* | OTHER | 0.972549 | 0.026941 | 0.000203 | 0.000247 | 0.000059 | |
| *Nostoc_sp._Q93TN0* | OTHER | 0.996702 | 0.002875 | 0.000405 | 0.00001 | 0.000008 | |

**Table S3A — Structural superposition of HMOX1 models. GmHMOX1 corresponds to PDB : 7cka reference enzymes.**

| Structural comparison | RMSD (Å) | Pruned atom pairs |
| --- | --- | --- |
| **GmHMOX1 vs CrHMOX1** | 0.557 | 183 |
| **GmHMOX1 vs PicoHMOX1** | 0.766 | 190 |
| **CrHMOX1 vs PicoHMOX1** | 0.607 | 179 |
| **PicoHMOX1 vs v1HMOX1** | 0.924 | 110 |
| **PicoHMOX1 vs v7HMOX1** | 1.025 | 121 |

**Table S3B — Structural superposition of PcyA models. SynPcyA corresponds to PDB : 2DE1 reference enzymes.**

| Structural comparison | RMSD (Å) | Pruned atom pairs |
| --- | --- | --- |
| **SynPcyA vs CrPcyA** | 1.018 | 186 |
| **SynPcyA vs PicoPcyA** | 0.987 | 195 |
| **CrPcyA vs PicoPcyA** | 0.784 | 233 |
| **PicoPcyA vs v3PcyA** | 0.893 | 176 |
| **PicoPcyA vs v8PcyA** | 0.936 | 135 |

*RMSD values correspond to the superposition after pruning of structurally divergent atom pairs using ChimeraX Matchmaker.*

**Table S4A — Conservation and divergence of structural features of HMOX1 models.**

| Feature | GmHMOX1 | PicoHMOX1 | Group I (v1–v5) | Group II (v7–v8) | Interpretation |
|---|---|---|---|---|---|
| **Overall HMOX1 fold** | Canonical | Conserved | Conserved | Conserved | Overall HMOX1 architecture retained |
| **Catalytic residues Gln34, Tyr144, Ala149, His150, Gly153** | Conserved | Conserved | Conserved | Conserved | Catalytic scaffold preserved |
| **Proximal His30** | Conserved | Conserved | Conserved | Conserved | Heme coordination preserved |
| **His30–heme Fe distance (Å)** | 1.936 Å | 1.989 Å | 2.120 Å (v1) | 2.060 Å (v7) | Consistent heme coordination |
| **Arg23 / Asn 145** | Conserved | Conserved | Conserved | **Absent** | Group-specific remodelling of propionate environment |
| **Heme-binding pocket** | Canonical | Canonical | Local rearrangement | Local rearrangement | Conserved catalytic architecture with local remodelling |
| **Heme Pocket entrance electrostatics** | Reference | Reference | More electropositive | More electropositive | Redistribution of surface charge |
| **Hydrophobicity** | Reference | Reference | Conserved | Conserved | Conserved hydrophobicity |
| **Structural interpretation** | Canonical HMOX1 | Canonical HMOX1 | Conserved catalytic scaffold with local remodelling | Conserved catalytic scaffold with group-specific remodelling | Group-specific surface properties |

**Table S4B — Conservation and divergence of structural features of PcyA models**

| Residues | **GmHMOX1** | **PicoHMOX1** | **Group I (v1-v5)** | **Group II (V7-V9)** | **Interpretation** |
|---|---|---|---|---|---|
| Glu76, His88, Asp105, Ser114, Arg149, Trp154, Asn219, Lys221, Thr222, and Phe244 | Conserved | Conserved | Conserved | Conserved | BV binding preserved |
| Val90 | Conserved | Conserved | Conserved | **Mutated in v7 and v8** | BV binding specificity of Group II vPcyA |
| Glu76, His88, Asp105, Tyr212, and Gln216 | Conserved | Conserved | Conserved | Conserved | Catalytic scaffold preserved |
| BV Pocket entrance electrostatics | Reference | Reference | Conserved | More electropositive | Redistribution of surface charge of group II vPcyA |
| Hydrophobicity | Reference | Reference | Conserved | Conserved | Conserved hydrophobicity |
